# SpaGE: Spatial Gene Enhancement using scRNA-seq

**DOI:** 10.1101/2020.05.08.084392

**Authors:** Tamim Abdelaal, Soufiane Mourragui, Ahmed Mahfouz, Marcel J.T. Reinders

## Abstract

Single-cell technologies are emerging fast due to their ability to unravel the heterogeneity of biological systems. While scRNA-seq is a powerful tool that measures whole-transcriptome expression of single cells, it lacks their spatial localization. Novel spatial transcriptomics methods do retain cells spatial information but can only measure tens to hundreds of transcripts. To resolve this discrepancy, we developed SpaGE, a method that integrates spatial and scRNA-seq datasets to predict whole-transcriptome expressions in their spatial configuration. Using five dataset-pairs, SpaGE outperformed previously published methods and showed scalability to large datasets. Moreover, SpaGE predicted new spatial gene patterns that are confirmed independently.

## Introduction

Single cell technologies rapidly developed over the last decade and have become valuable tools for enhancing our understanding of biological systems. Single-cell RNA-sequencing (scRNA-seq) allows unbiased measurement of the entire gene expression profile of each individual cell and has become the de facto technology used to characterize the cellular composition of complex tissues [1, 2]. However, single cells often have to be dissociated before performing scRNA-seq and results in losing the spatial context and hence limits our understanding of cell identities and relationships. Recently, spatial transcriptomics technologies have advanced and provide localizations of gene expressions and cellular structure at the cellular level [3, 4]. Current protocols can be divided in two categories: 1) imaging-based methods (e.g. osmFISH and MERFISH) [5, 6], and 2) sequencing-based methods (e.g. STARmap and Slide-seq) [7, 8]. Imaging-based protocols have a high gene detection sensitivity; capturing high proportion of the mRNA molecules with relatively small dropout rate. However, imaging-based protocols are often limited in the number of genes that can be measured simultaneously. On the other hand, sequencing-based protocols like STARmap can scale up to thousands of genes, it has a relatively lower gene detection sensitivity. Slide-seq is not limited in the number of measured genes and can be used to measure the whole transcriptome. However, similar to STARmap, Slide-seq suffers from a low gene detection sensitivity. In addition, osmFISH, MERFISH and STARmap can capture genes at the single-molecule resolution, which can be averaged or aggregated to the single-cell level. While Slide-seq has a resolution of 10μm, which is comparable to the average cell size, but does not always represent a single-cell.

Given the complementary information provided by both scRNA-seq and spatial transcriptomics data, integrating both types would provide a more complete overview of cell identities and interactions within complex tissues. This integration can be performed in two different ways [9]: 1) dissociated single-cells measured with scRNA-seq can be mapped to their physical locations in the tissue [10–12], or 2) missing gene expression measurements in the spatial data can be predicted from scRNA-seq. In this study, we focus on the second challenge in which measured gene expressions of spatial cells can be enhanced by predicting the expression of unmeasured genes based on scRNA-seq data of a matching tissue. Several methods have addressed this problem using various data integration approaches to account for the differences between the two data types [13–16]. All these methods rely on joint dimensionality reduction methods to embed both spatial and scRNA-seq data into a common latent space. For example, Seurat uses canonical correlation analysis (CCA), Liger uses non-negative matrix factorization (NMF), and Harmony uses principal component analysis (PCA). While Seurat, Liger and Harmony rely on linear methods to embed the data, gimVI uses a non-linear deep generative model. Despite recent benchmarking efforts [17], a comprehensive evaluation of these methods for the task of spatial gene prediction from dissociated cells is currently lacking. For example, Seurat, Liger and gimVI, have only been tested using relatively small datasets (<2,000 cells) [13, 14, 16]. It is thus not clear whether a complex model, like gimVI, is really necessary. Moreover, Seurat, Harmony and gimVI lack interpretability of the integration procedure, so that it does not become clear which genes contribute in the prediction task.

Here, we present SpaGE (Spatial Gene Enhancement), a robust, scalable and interpretable machine-learning method to predict unmeasured genes of each cell in spatial transcriptomic data through integration with scRNA-seq data from the same tissue. SpaGE relies on domain adaptation using PRECISE [18] to correct for differences in sensitivity of transcript detection between both single-cell technologies, followed by a k-nearest-neighbor (kNN) prediction of new spatial gene expression. We demonstrate that SpaGE outperforms state-of-the-art methods by accurately predicting unmeasured gene expression profiles across a variety of spatial and scRNA-seq dataset pairs of different regions in the mouse brain. These datasets include a large spatial data with more than 60,000 cells, used to illustrate the scalability and computational efficiency of SpaGE compared to other methods.

## Results

### SpaGE overview

We developed SpaGE, a platform that enhances the spatial transcriptomics data by predicting the expression of unmeasured genes, from a dissociated scRNA-seq data from the same tissue (Fig. 1). Based on the set of shared genes, we align both datasets using the domain adaptation method PRECISE [18], to account for technical differences as well as gene detection sensitivity differences. PRECISE geometrically aligns linear latent factors computed on each dataset and finds gene combinations expressed in both datasets. These gene combinations thus define a common latent space and can be used to jointly project both datasets. Next, in this common latent space, we use the kNN algorithm to define the neighborhood of each cell in the spatial data from the scRNA-seq cells. These neighboring scRNA-seq cells are then used to predict the expression of spatially unmeasured genes. Finally, we end up with the full gene expression profile of each cell in the spatial data.

**Fig. 1.**
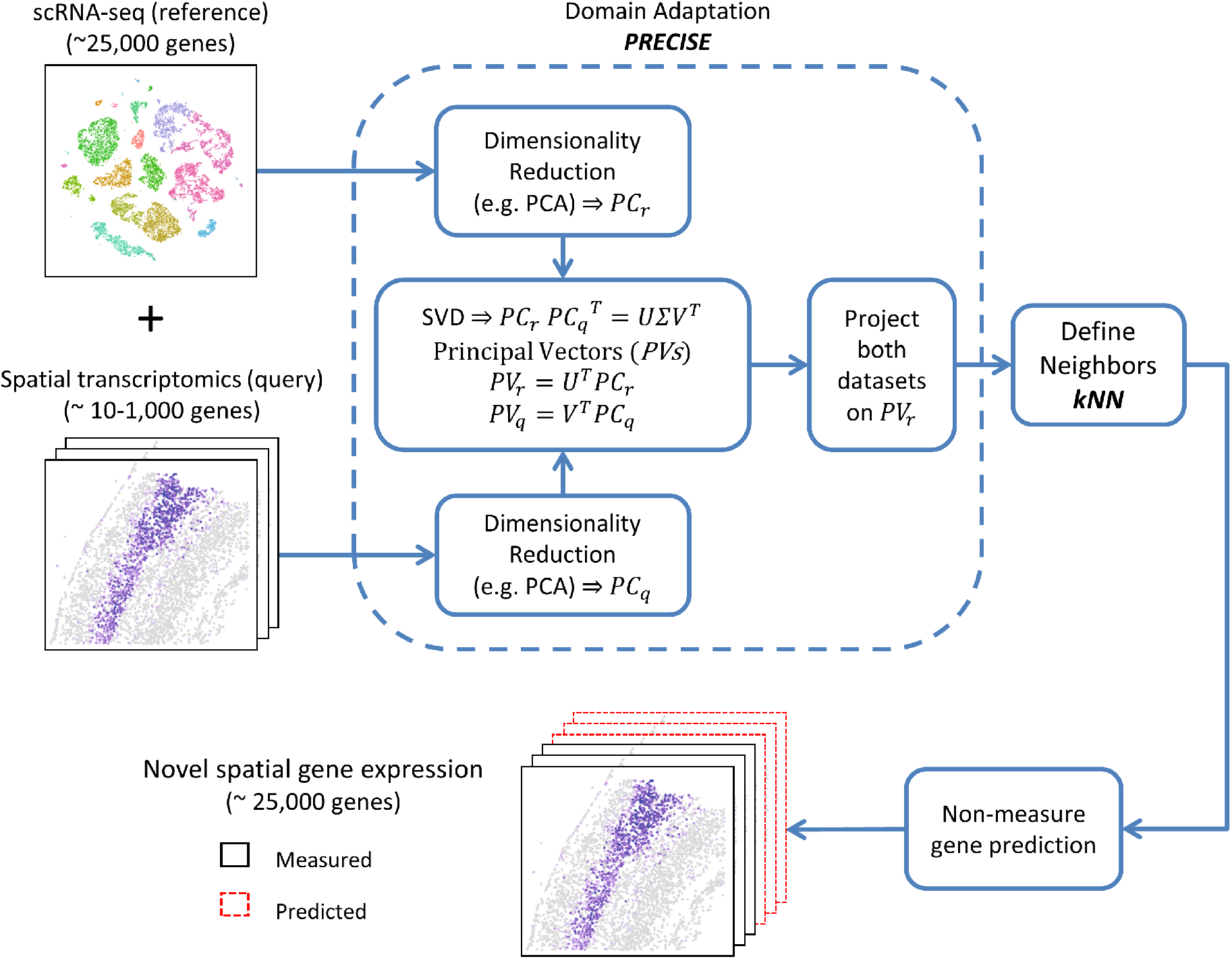
SpaGE pipeline. SpaGE takes as input two datasets, a scRNA-seq dataset and a spatial transcriptomics dataset measured from the same tissue. SpaGE uses gene combinations of equal significance in both datasets to predict spatial locations of unmeasured genes. Using PRECISE, SpaGE finds directions that are important for both datasets, by making use of a geometrical alignment of the independent *PCs* to produce the *PVs*. SpaGE aligns both datasets by projecting on the *PVs* of the reference dataset. Using the aligned datasets, SpaGE applies kNN prediction to define new gene expression patterns for spatially unmeasured genes, predicted from the dissociated scRNA-seq data. Each spatial cell can be enhanced by having the expression of the whole transcriptome.

The alignment step is the most crucial step in the pipeline of SpaGE. For this purpose, we use PRECISE, a domain adaptation method previously proposed to predict the drug response of human tumors based on pre-clinical model such as cell lines and mouse models. We adapted PRECISE to the task of integrating the spatial data with the scRNA-seq data by defining the common aligned subspace between both datasets (Fig. 1). PRECISE takes as input the expression matrix of both datasets, having the same set of (overlapping) genes but measured differently and within different cells. As we are aiming to fit each spatial cell to the most similar scRNA-seq cells, we may refer to the spatial dataset as the ‘query’ and the scRNA-seq dataset as the ‘reference’. First, PRECISE obtains a lower dimensional space for each dataset separately using a linear dimensionality reduction method, such as Principal Component Analysis (PCA). Next, the two independent sets of principal components (*PCs*) are aligned by applying a singular value decomposition. We align the two sets of principal components using the singular vectors to obtain the aligned components, named principal vectors (*PVs*). These *PVs* are sorted in decreasing order based on their similarity between the reference and the query datasets. This allows us to filter out dissimilar or noisy signals, by discarding *PVs* with relatively low similarity, thus keeping only the common latent space. The principal vectors of the reference dataset (*PV_r_*) are considered as the aligned latent space. We project both datasets on *PV_r_* to obtain the new aligned versions used for the kNN prediction.

We performed SpaGE on five dataset pairs from different regions in the mouse brain, varying in the number of cells and the number of spatially measured genes, summarized in Table 1. To show the alignment performance, we calculated the cosine similarity between the *PCs* and the *PVs* i.e. before and after the alignment. Across all five dataset pairs, we observed that indeed the relation between the *PCs* is not one-to-one, as these *PCs* are obtained from two different datasets (Figure S1-2). However, after alignment using PRECISE, the diagonal cosine similarity between the *PVs* is maximized showing a one-to-one relationship between the *PVs* of both datasets. Figure S1A shows the diagonal cosine similarity before and after PRECISE (i.e. between *PCs* and *PVs*) across all dataset pairs, showing a relatively large increase in similarity after the alignment using PRECISE.

**Table 1.** Summary of the dataset pairs used in this study.

| Spatial_scRNA-seq dataset pair | Spatial data |  |  | scRNA-seq data |  |  |
| --- | --- | --- | --- | --- | --- | --- |
|  | # of cells | # of genes | Tissue | # of cells | # of genes | Tissue |
| STARmap_AllenVISp [7, 19] | 1,549 | 1,020 | Visual cortex | 14,249 | 34,617 | Visual cortex |
| osmFISH_Zeisel [5, 20] | 3,405 | 33 | Somatosensory cortex | 1,691 | 15,075 | Somatosensory cortex |
| osmFISH_AllenSSp [5, 21] |  |  |  | 5,613 | 30,527 |  |
| osmFISH_AllenVISp [5, 19] |  |  |  | 14,249 | 34,617 | Visual cortex |
| MERFISH_Moffit [4] | 64,373 | 155 | Pre-optic region | 31,299 | 18,646 | Pre-optic region |

Another interesting feature of SpaGE is the ability to interpret the most contributing genes defining the latent integration space. In general, these genes are highly variable and in most cases are related to cell type differences. A good example is the integration of the **osmFISH_Zeisel** dataset pair, in which the top six contributing genes are *Tmem2, Mrc1, Kcnip2, Foxj1, Apln* and *Syt6* (Methods). These genes are related to six different cell categories previously defined in the **osmFISH** paper [5]: Oligodendrocytes, Immune cells, Inhibitory neurons, Ventricle, Vasculature and Excitatory neurons, respectively.

### SpaGE outperforms state-of-the-art methods on the STARmap dataset

Using the first dataset pair **STARmap_AllenVISp**, we applied SpaGE to integrate both datasets and predict unmeasured spatial gene expression patterns. In order to evaluate the prediction, we performed a leave-one-gene-out cross validation (Methods). The **STARmap_AllenVISp** dataset pair shares 994 genes. In each crossvalidation fold, one gene is left out and the remaining 993 genes are used as input for SpaGE to predict the spatial expression pattern of the left-out gene. We evaluated the prediction performance by calculating the Spearman correlation between the original measured spatially distributed values and the predicted values of the left-out gene. We performed the same leave-one-gene-out cross validation using Seurat, Liger and gimVI, to benchmark the performance of SpaGE. Results show a significant improvement in performance for SpaGE compared to all three methods (p-value <0.05, two-sided paired Wilcoxon rank sum test), with a median Spearman correlation of 0.125 compared to 0.083, 0.067 and 0.055 for Seurat, Liger and gimVI, respectively (Fig. 2a).

**Fig. 2.**
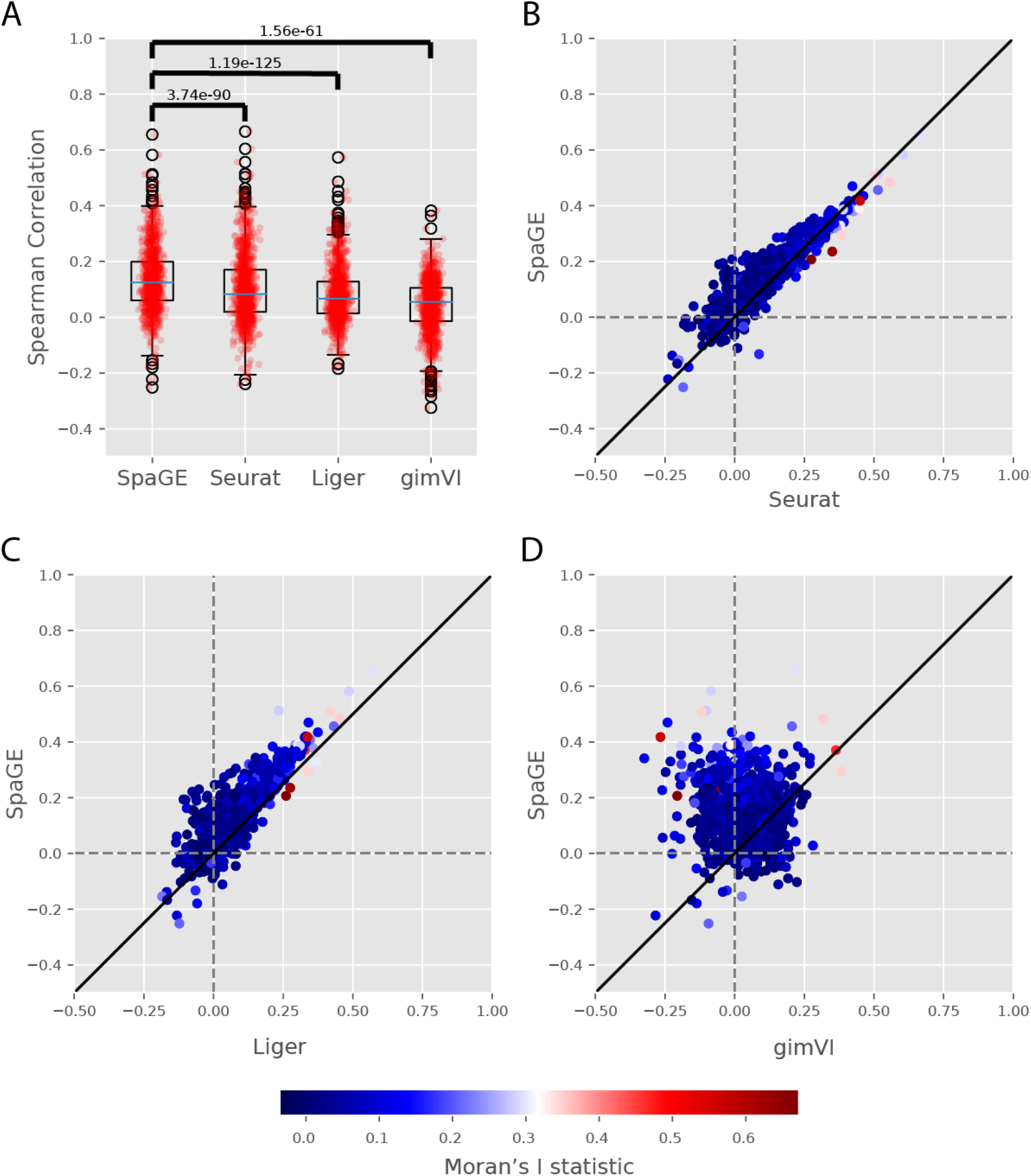
Prediction performance comparison for the STARmap_AllenVISp dataset pair. **(A)** Boxplots showing the Spearman correlations for the leave-one-gene-out cross validation experiment for each method. The blue lines show the median correlation across all genes with a better performance for SpaGE. The red dots show the correlation values for individual genes. The p-values show the significant difference between all correlation values of SpaGE and each other method, using a paired Wilcoxon rank-sum test. **(B-D)** Detailed performance comparison between SpaGE and **(B)** Seurat, **(C)** Liger, **(D)** gimVI. These scatter plots show the correlation value of each gene across two methods. The solid black line is the y=x line, the dashed lines show the zero correlation. Points are colored according to the Moran’s I statistic of each gene. All scatter plots show that the majority of the genes are skewed above the y=x line, showing an overall better performance of SpaGE over other methods.

Further, we compared the Spearman correlation of SpaGE versus the state-of-the-art methods per gene, to obtain a detailed evaluation. Results show better performance of SpaGE across the majority of genes, but not all (Fig. 2b-d). Next, we visually compared a few genes that had high correlations for each method. For the top three predicted genes of SpaGE *(Pcsk2, Pgm2l1* and *Egr1*), Seurat obtained a good prediction as well, as these three genes are in the top 10 predicted genes of Seurat. Liger failed to predict *Egr1*, while gimVI failed to predict *Pgm2l1* and *Egr1* (Figure S3A). We further looked for examples where other methods obtained higher correlations than SpaGE, excluding the top 10 predicted genes by SpaGE. Compared to Seurat, SpaGE similarly predicted the expression of *Arpp19*, but predicted relatively higher contrast patterns for *Pcp4* and *Arc* (Figure S3B). Compared to Liger, SpaGE similarly predicted the expression of *Mobp,* higher contrast pattern for *Hpcal4,* and better predicted the spatial pattern of *Tsnax* (Figure S3C). Compared to gimVI, SpaGE obtained good prediction for *Tsnax,* predicted wrong pattern for *Ythdc2,* and lower contrast prediction for *Bcl6* (Figure S3D).

Although the correlation values are in general low, SpaGE is capable of accurately reconstructing genes with clear spatial pattern in the brain. Fig. 3 shows a set of genes known to have spatial patterns (previously reported by Seurat, Liger and gimVI). In this set of genes, Seurat and Liger are performing well, except that Liger produced a lower contrast expression pattern in some cases (e.g. *Lamp5* and *Bsg*). However, gimVI is unable to predict the correct gene pattern, and even predicts a reverse pattern in the case of *Plp1*.

**Fig. 3.**
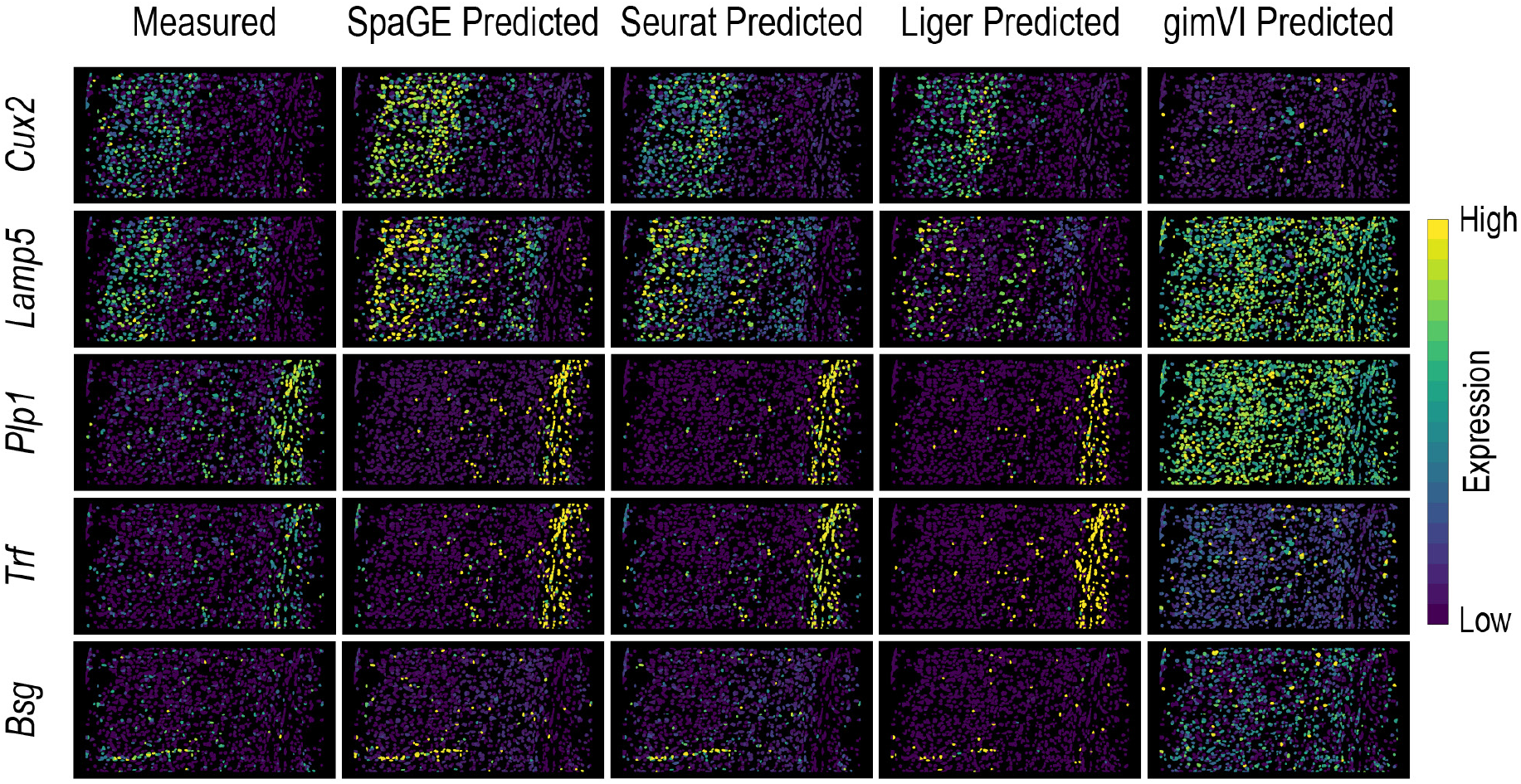
Predicted expression of known spatially patterned genes in the STARmap dataset. Each row corresponds to a single gene having a clear spatial pattern. First column from the left shows the measured spatial gene expression in the **STARmap** dataset, while other columns show the corresponding predicted expression pattern by SpaGE, Seurat, Liger and gimVI, using the leave-one-gene-out cross validation experiment. Prediction is performed using the **AllenVISp** dataset.

Additionally, although it is important to accurately predict the expression of all genes, genes with distinct spatial patterns are more important to accurately predict compared to non-or randomly expressed genes. To quantify the existence of spatial patterns, we calculate the Moran’s I statistics of each gene using the original **STARmap** spatial data (Methods). We compared the prediction performance of each gene with the corresponding Moran’s I value. For SpaGE, Seurat and Liger, we observed a positive relationship between the prediction performance and the Moran’s I values, i.e. genes with spatial patterns are better predicted (Figure S4A-C). On the other hand, gimVI performed worse on genes with high Moran’s I statistics (Figure S4D).

In addition, we evaluated how well a gene can be predicted when using less shared genes. First, we selected a fixed test set of 50 genes, next we down-sampled the remaining set of 944 shared genes in a guided manner (Methods). For downsampled shared genes sets of 10, 30, 50, 100, 200 and all 944 genes, SpaGE performance always increases with the number of shared genes as expected (Figure S4E).

### SpaGE predicts unmeasured spatial gene patterns that are independently validated

After validating SpaGE to accurately predict the spatially measured genes, we applied SpaGE to predict new unmeasured genes for the spatial data, with the aim to define novel spatial gene patterns. We illustrate SpaGE’s capability of such task using the **STARmap_AllenVISp** dataset pair. First, during the leave-one-gene-out cross validation, SpaGE was able to produce the correct spatial pattern for *Rorb*, *Syt6* and *Tbr1* (Fig. 4). These three genes were originally under-expressed, possibly due technical noise or low gene detection sensitivity in the **STARmap** dataset. Our predictions using SpaGE are in agreement with the highly sensitive cyclic smFISH dataset (**osmFISH** [5]) measured from the mouse somatosensory cortex, a similar brain region in terms of layering structure to the visual cortex measured by the **STARmap** dataset. Further, using SpaGE, we were able to obtain novel spatial gene patterns for five genes not originally measured by the **STARmap** dataset (Fig. 5). These predicted patterns are supported by the Allen Brain Atlas in-situ hybridization (Allen ISH).

**Fig. 4.**
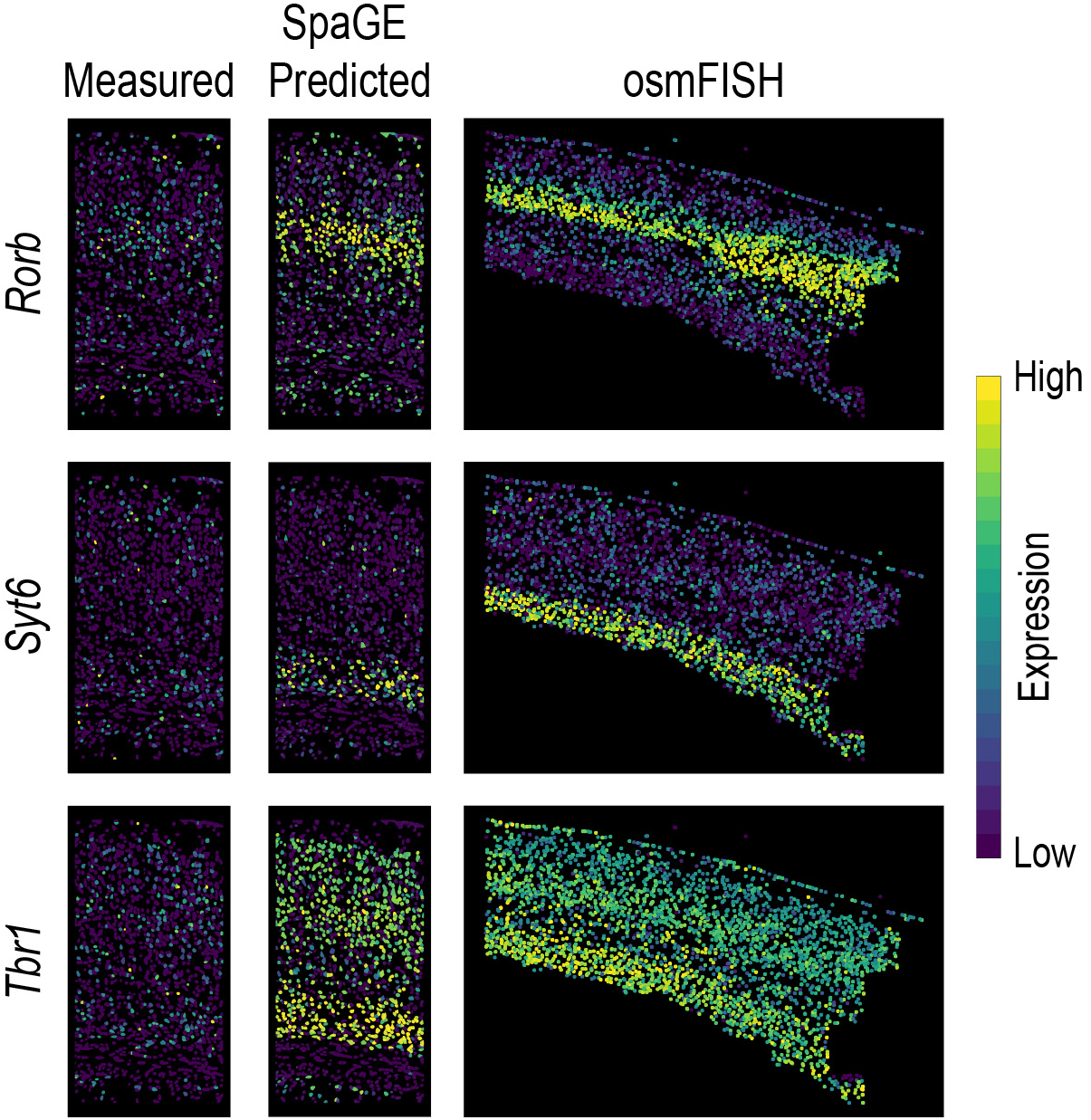
SpaGE accurately predicted the expression of *Rorb, Syt6* and *Tbr1* in agreement with the osmFISH data. These three genes (shown in rows) were wrongly measured in the original **STARmap** data (shown in the left column). Using the **STARmap**_AllenVISp dataset pair, SpaGE was able to reconstruct the correct spatial gene expression patterns (middle column). These predicted patterns are in agreement with the measured gene expression patterns measure by the **osmFISH** dataset (right column), a highly sensitive single-molecule technology.

**Fig. 5.**
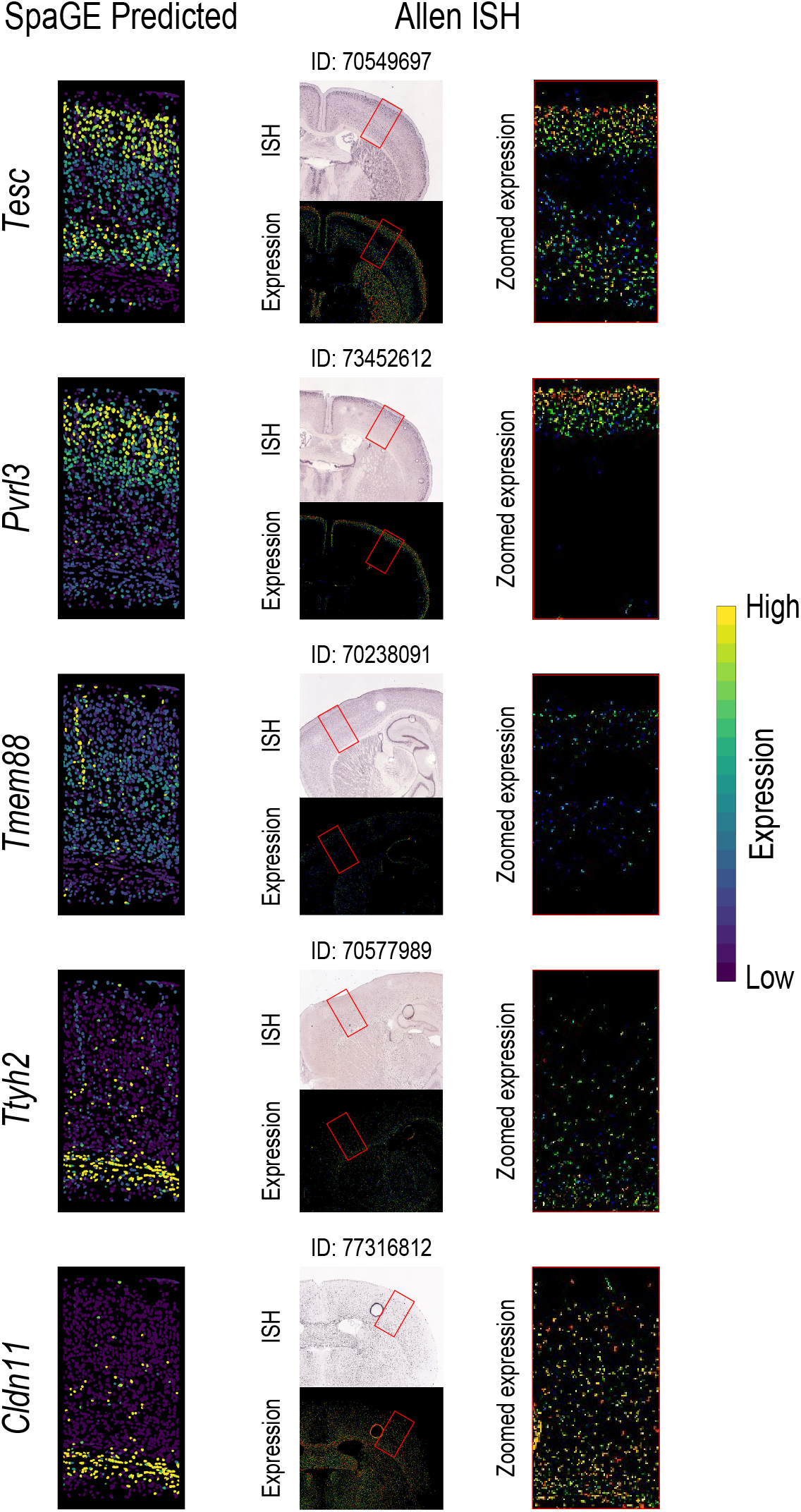
Novel gene expression patterns for five genes not originally measured by the STARmap dataset, validated using the Allen Brain Atlas in-situ hybridization ISH. The left column shows the predicted spatial patterns using SpaGE for these five genes (shown in rows). The middle column shows the Allen ISH data for each gene, stating the image ID on top of each tissue section. The red rectangle highlights the corresponding brain region measured by the **STARmap** dataset. The right column shows a zoomed-in view of the region highlighted using this red rectangle, showing an agreement with the expression patterns predicted by SpaGE.

### SpaGE produces better prediction with deeply sequenced reference dataset

We wanted to test the effect of changing the reference scRNA-seq data on the spatial gene expression prediction. Here, we used the **osmFISH** dataset which represents a different challenge compared to the **STARmap** dataset. On one hand, the **osmFISH** dataset has a relatively higher gene detection sensitivity, but on the other hand, the **osmFISH** dataset includes only 33 genes. First, we evaluated the **osmFISH_Zeisel** dataset pair, in which we integrated the **osmFISH** dataset with a reference scRNA-seq dataset from the same lab (**Zeisel** et. al [20]). We performed leave-one-gene-out cross validation similar to the **STARmap** dataset. Compared to other methods, SpaGE has significantly better performance (p-value <0.05, two-sided paired Wilcoxon rank sum test), with a median Spearman correlation of 0.203 compared to 0.007, 0.090 and 0.130 for Seurat, Liger and gimVI, respectively (Fig. 6, Figure S5A). For a more detailed comparison per gene: SpaGE is performing better on the majority of genes compared to Liger and gimVI, while compared to Seurat, SpaGE has better performance across all genes (Figure S5B-D). We further investigated the relation between the prediction performance and the Moran’s I statistics of the originally measured genes. Similar to the STARmap data, for SpaGE and Seurat, we found a positive relationship, i.e. the performance is higher for genes with distinct spatial patterns. However, Liger and gimVI have a negative relationship (Figure S6).

**Fig. 6.**
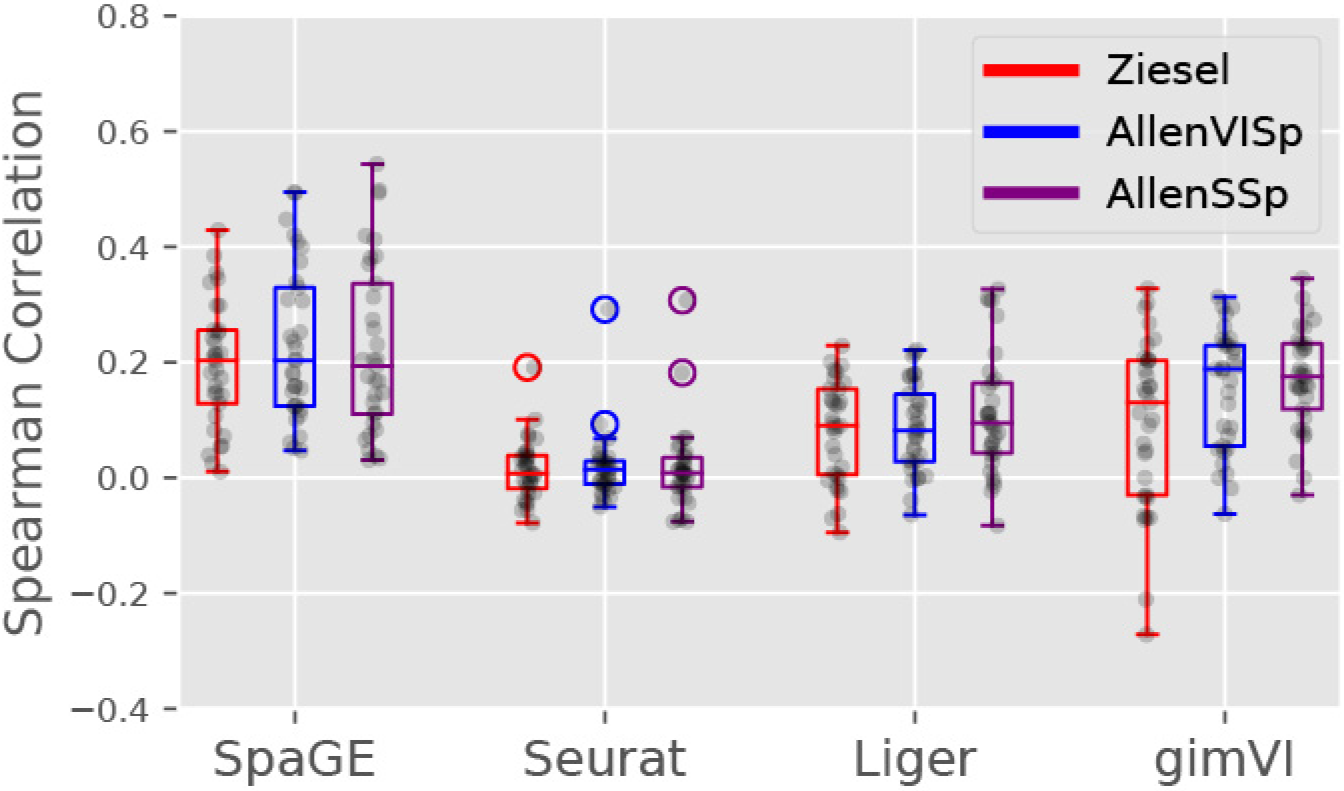
Prediction performance comparison for the osmFISH dataset using different reference scRNA-seq datasets. Boxplots showing the Spearman correlations for the leave-one-gene-out cross validation experiment for each method using three different scRNA-seq datasets, **Zeisel**, **AllenVISp** and **AllenSSp**. The median correlations shows a better performance for SpaGE in all dataset pairs. The black dots show the correlation values for individual genes. SpaGE showed a performance improvement when using the **AllenVISp** over the **Zeisel** data. Although the median correlation is the same, the overall correlation range did improve. Also, gimVI clearly benefits from using the **AllenVISp** and the **AllenSSp** datasets over the **Zeisel** dataset.

Next, we tested the performance of all methods using the **AllenVISp** dataset as reference for the **osmFISH** dataset, similar to the **STARmap** dataset. For the **osmFISH_AllenVISp** dataset pair, we observed similar conclusions where SpaGE has significantly better performance compared to other methods, with a median Spearman correlation of 0.203 compared to 0.014, 0.082 and 0.189 for Seurat, Liger and gimVI, respectively (Fig. 6, Figure S7A). SpaGE has better performance across all genes compared to Seurat and Liger, while gimVI is performing better on a few genes (Figure S7B-D). All four methods have a positive relationship between their prediction performance and the Moran’s I statistics of the measured genes (Figure S8). These results show how the reference dataset can affect the prediction. Compared to the **Zeisel** dataset, the **AllenVISp** is more deeply sequenced data, with the average number of detected transcripts per cell being ~140x more than the **Zeisel** dataset (Figure S9A-B). However, not all methods benefit from this, as for Seurat and Liger, the prediction performance using the **AllenVISp** or the **Zeisel** datasets is quite similar (Fig. 6). On the other hand, SpaGE and gimVI get an increase in performance across all genes, although the median correlation for SpaGE remains the same.

While the **AllenVISp** is a deeply sequenced reference dataset, it has been measured from a different brain region than the **osmFISH** dataset (Table 1). Therefore, we decided to use a third reference dataset, **AllenSSp**, which has roughly the same sequencing depth as the **AllenVISp** (Figure S9B-C) but is measured from the somatosensory cortex, similar to the **osmFISH** dataset. We evaluated the prediction performance of all four tools for the new dataset pair **osmFISH_AllenSSp**. Again, SpaGE obtained a better performance with a median Spearman correlation of 0.194 compared to 0.008, 0.095 and 0.176 for Seurat, Liger and gimVI, respectively (Fig. 6, Figure S10A). SpaGE has a better performance across all genes compared to Seurat, and better performance across the majority of genes compared to Liger and gimVI (Figure S10B-D). Similar to previous observations, SpaGE, Liger and gimVI have positive relationship between the prediction performance and Moran’s I statistics. However, Seurat has a negative relationship (Figure S11).

Changing the brain region did not affect the overall performance of SpaGE (Fig. 6), however, the prediction of genes with known patterns did improve (Fig. 7). When we visually inspect these genes, we can clearly observe that the predicted spatial pattern improved when the sequencing depth of the reference set improves or becomes from a similar tissue. *Rorb* and *Tbr1* are clear examples, where the prediction using **Zeisel** was almost missing the correct pattern, this became clearer using the **AllenVISp** having a greater sequencing depth. Second, changing to a matching tissue adds further improves the predicted patterns of these genes (**AllenSSp**). Eventually, all five genes *(Lamp5, Kcnip2, Rorb, Tbr1* and *Syt6)* are more accurately predicted using the **AllenSSp** dataset.

**Fig. 7.**
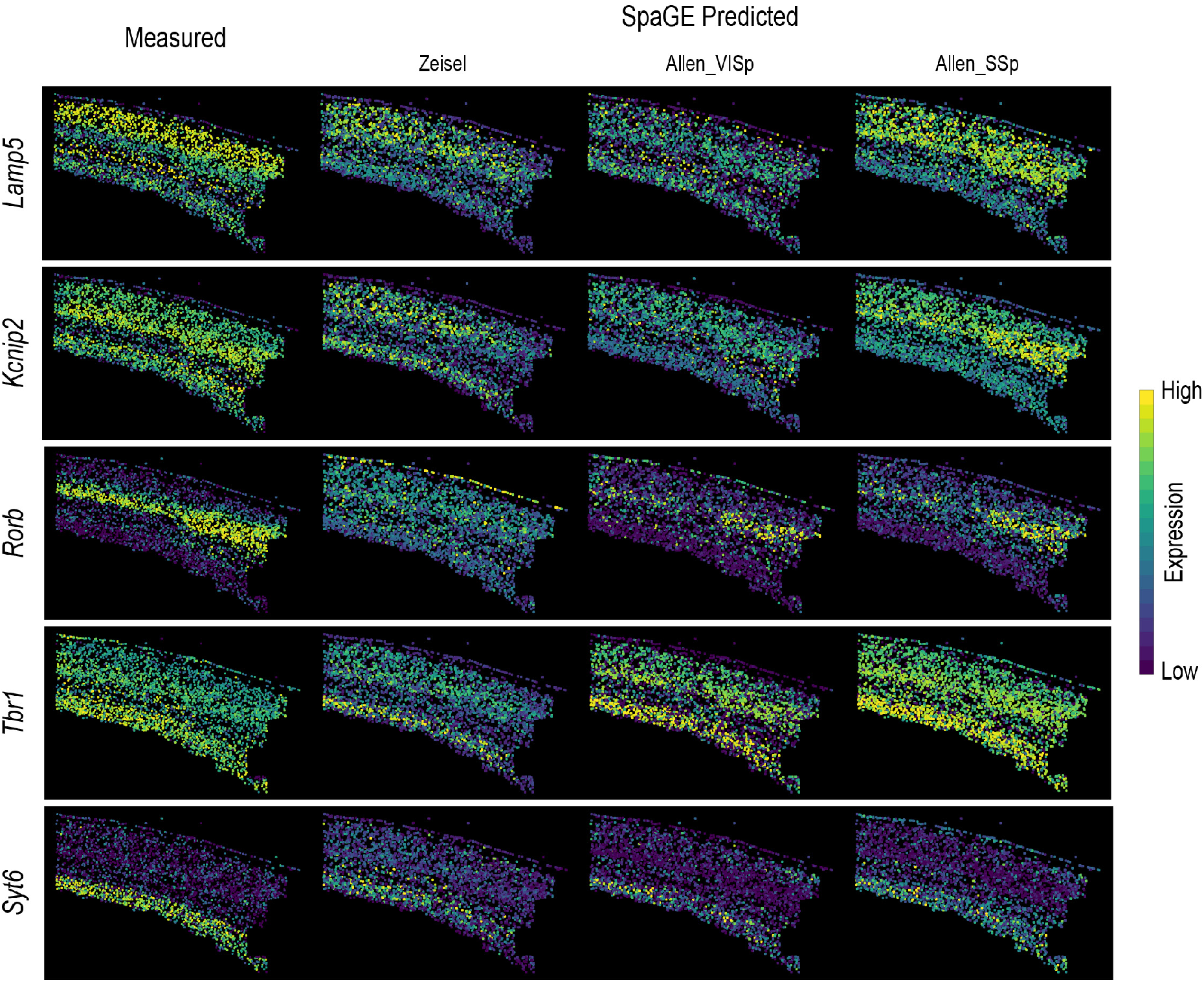
Predicted expression of known spatially patterned genes in the osmFISH dataset using different reference scRNA-seq datasets. Each row corresponds to a single gene having a clear spatial pattern. First column from the left shows the measured spatial gene expression in the **osmFISH** dataset, while the second, third and fourth columns show the corresponding predicted expression pattern by SpaGE using **Zeisel**, **AllenVISp** and **AllenSSp** datasets, respectively. Changing from **Zeisel** to **AllenVISp** (deeply sequenced data) improved the prediction, while matching the brain region using the **AllenSSp** improved the prediction further.

### SpaGE is scalable to large spatial datasets

So far, SpaGE showed good prediction performance in the leave-one-gene-out predictions, and was also able to predict correct spatial patterns of unmeasured genes within the spatial transcriptomic datasets. All these results were, however, obtained using a relatively small spatial datasets including only a few thousand cells (**STARmap** and **osmFISH**). This opens the question of how does SpaGE scale to large spatial datasets, comparable to the datasets measured nowadays. To assess the scalability of SpaGE, we used a large **MERFISH** dataset with >60,000 cells measured from the mouse brain pre-optic region, and integrated it with the corresponding scRNA-seq dataset published in the same study by **Moffit** et al [4]. The **MERFISH_Moffit** dataset pair shares 153 genes on which we applied the same leave-one-gene-out cross validation using all four methods. Similar to the previous results, SpaGE significantly outperformed all other methods (p-value <0.05, two-sided paired Wilcoxon rank sum test) with a median Spearman correlation of 0.275 compared to 0.258, 0.027 and 0.099 for Seurat, Liger and gimVI, respectively (Fig. 8A). Per gene comparisons shows a clear advantage of SpaGE versus Liger and gimVI, but more comparable performance with Seurat (Fig. 8B-D).

**Fig. 8.**
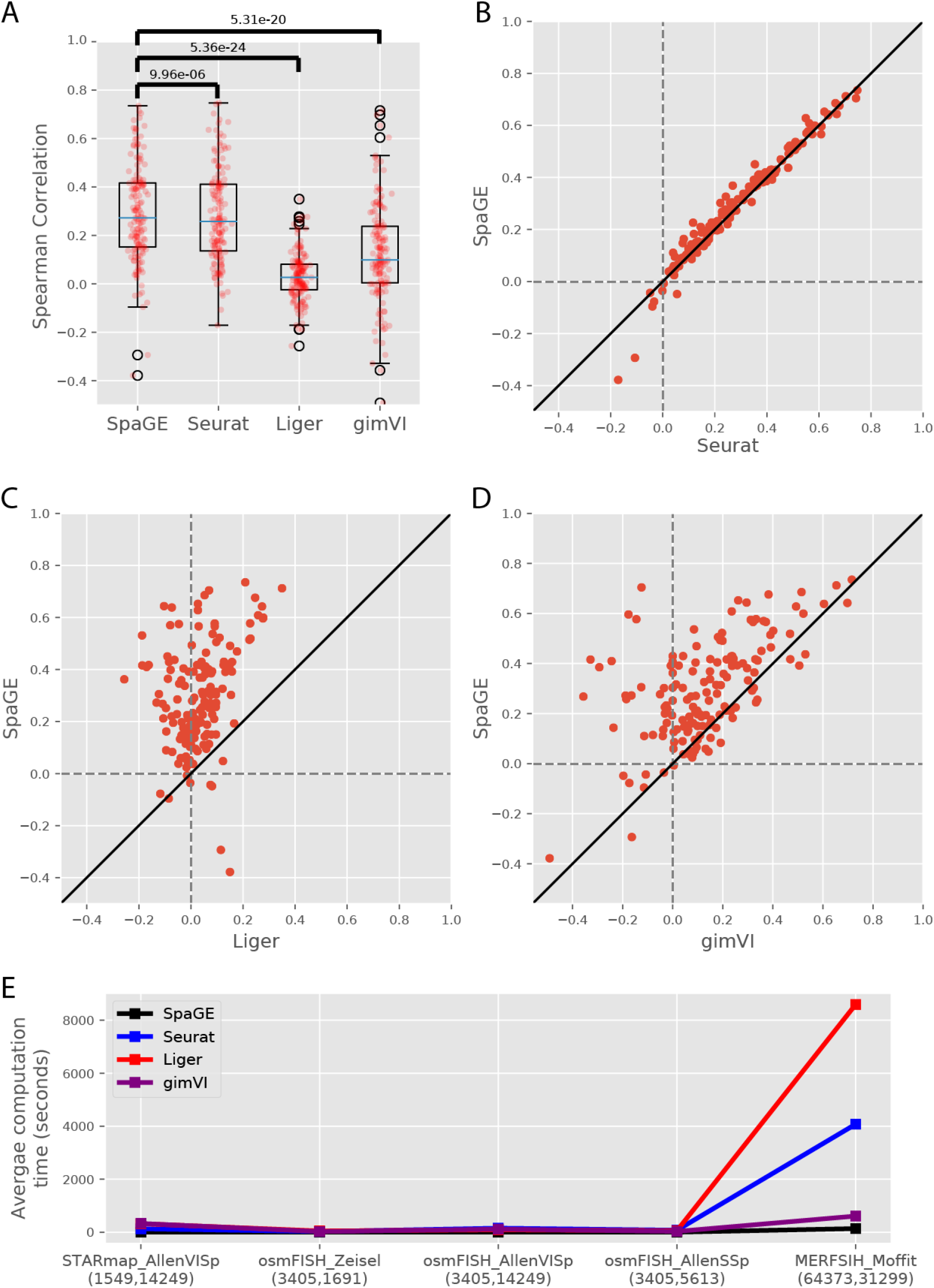
Prediction performance comparison for the MERFISH_Moffit dataset pair. **(A)** Boxplots showing the Spearman correlations for the leave-one-gene-out cross validation experiment for each method. The blue lines show the median correlation across all genes with a better performance forSpaGE. The red dots show the correlation values for individual genes. The p-values show the significant difference between all correlation values of SpaGE and each other method, using a paired Wilcoxon rank-sum test. **(B-D)** Detailed performance comparison between SpaGE and **(B)** Seurat, **(C)** Liger, **(D)** gimVI. These scatter plots show the correlation value of each gene across two methods. The solid black line is the y=x line, the dashed lines show the zero correlation. All scatter plots show that the majority of the genes are skewed above the y=x line, showing an overall better performance of SpaGE over other methods. **(E)** Line plots showing the average computation time per gene of each method for each dataset pair. Numbers below each dataset pair show the number of cells in the spatial and the scRNA-seq datasets, respectively.

Further, we compared the computation times of all four methods across all five dataset pairs. All experiments were run on a Linux HPC server but limited to a single CPU core, with 256 GB of memory, to be able to compare runtimes. The calculated computation time includes the integration and the prediction time. We did not observe large differences between the methods when testing for the **STARmap** and the **osmFISH** datasets, as these datasets are relatively small (Fig. 8E), even though SpaGE has the lowest average computation time per gene. For the large MERFISH dataset, SpaGE has a clear advantage compared to the other methods as the average computation time of SpaGE is ~30x, 63x and 4x faster than Seurat, Liger and gimVI, respectively. These results combined show an overall advantage of SpaGE over other methods for larger datasets with higher prediction performance and lower computation time.

## Discussion

We demonstrated the ability of SpaGE to enhance spatial transcriptomics data by predicting the expression of unmeasured genes based on scRNA-seq data collected from the same tissue. The ability of SpaGE to produce accurate gene expression prediction highly depends on the alignment part performed using PRECISE, which rotates the principal components of each dataset to produce principal vectors with high one-to-one similarity. Projecting the datasets to the latent space spanned by these principal vectors produces a proper alignment, making a simple kNN prediction sufficient to achieve accurate gene expression estimation.

We benchmarked SpaGE against three state-of-the-art methods for multi-omics data integration, using five different dataset pairs. These dataset pairs represent different challenges to the integration and prediction task, as they differ in gene detection sensitivity level and the number of spatially measured genes, which are the basis for the alignment. Increasing the number of shared genes should, in principle, eases the integration task and produces better prediction of unmeasured genes. Further, imaging-based spatial transcriptomic methods, with high gene detection sensitivity, may also improve the integration and prediction, as they are able to capture the majority of the genes even the ones with relatively low expression. On the other hand, integrating this high sensitivity data with scRNA-seq, which has lower sensitivity, can be more challenging. That is because the differences in gene expression are higher compared to integrating a sequencing-based spatial data with scRNA-seq data, both having comparable sensitivity.

Across all tested dataset pairs, SpaGE outperformed all methods producing better predictions for the majority of the genes. However, for few genes, SpaGE had lower prediction performance than other methods. Seurat produced good gene predictions for the **STARmap** and the **MERFISH** datasets, with similar predictions to SpaGE. However, Seurat did not properly work when there are very few shared genes, such as in the **osmFISH** dataset (33 genes). This problem is even more pronounced for Liger, as it performed relatively well for the **STARmap** dataset producing good gene predictions, but has a decreased performance for both the **osmFISH** (33 genes) and the **MERFISH** (155 genes) datasets. On the other hand, gimVI performed relatively well for the **osmFISH** and the **MERFISH** datasets. However, gimVI had overall the lowest performance for the **STARmap** dataset, with inaccurate predictions for genes with spatial patterns such as *Cux2* and *Lamp5*, and good predictions for lowly expressed or non-patterned genes such as *Ythdc2* and *Bcl6*. This suggests that gimVI works well with imaging-based technologies having high gene detection sensitivity, but not with the sequencing-based technologies.

Next to the overall best performance, SpaGE is an interpretable algorithm as it allows to find the genes driving the correspondence between the datasets. The principal vectors, used to align the datasets to a latent space, show the contribution of each gene in defining this new latent space. Further, SpaGE is scalable to large spatial data with significantly lower computation time compared to the other methods, as shown on the **MERFISH** dataset having more than 60,000 cells measured spatially. Moreover, SpaGE is a flexible pipeline. Here we used PCA as the initial independent dimensionality reduction algorithm. However, this step can be replaced by any linear dimensionality reduction method.

We used the Spearman Rank correlation to quantitatively evaluate the predicted gene expressions. The overall evaluation showed relatively low correlations across all methods and all dataset pairs. These low correlations express the difficulty of the problem, as the predicted gene expressions are obtained from a different type of data. However, the Spearman correlation is not the optimal evaluation metric, as it does not always reflect the spatially predicted patterns, i.e. visual inspection showed good predictions for genes with known spatial pattern in the mouse cortex, while the correlation values were less than 0.2.

## Conclusion

SpaGE presents a robust, scalable, interpretable and flexible method for predicting spatial gene expression patterns. SpaGE uses domain adaptation to align the spatial transcriptomics and the scRNA-seq datasets to a common space, in which unmeasured spatial gene expressions can be predicted. SpaGE is less complex and much faster when compared to other approaches and generalizes better across datasets and technologies.

## Methods

### SpaGE algorithm

The SpaGE algorithm takes as input two gene expression matrices corresponding to the scRNA-seq data (reference) and the spatial transcriptomics data (query). Based on the set of shared genes between the two datasets, SpaGE enriches the spatial transcriptomics data using the scRNA-seq data, by predicting the expression of spatially unmeasured genes. The SpaGE algorithm can be divided in two major steps: 1) Alignment of the two datasets using the domain adaptation algorithm PRECISE [18], and 2) gene expression prediction using k-nearest-neighbor regression.

First, PRECISE was used to project both datasets into a common latent space. Let *R*_(*n × g*)_ be the gene expression matrix of the reference dataset having *n* cells and *g* genes, and let *Q*_(*m × h*)_ be the gene expression matrix of the query dataset having *m* cells and *h* genes. Using the set of shared genes *p = g* ⋂ *h,* PRECISE applies independent Principal Component Analysis (PCA) for each dataset to define two independent sets of Principal Components (PCs), such that:

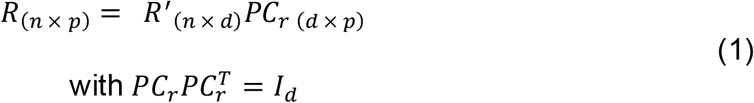

and

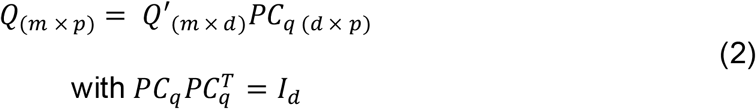

where *d* is the number of desired PCs, *PC_r_* and *PC_q_* represents the principal components of the reference and the query datasets, respectively. We choose *d* = 50 for the **STARmap_AllenVISp** and **MERFISH_Moffit** dataset pairs, and *d* = 30 for all the **osmFISH** dataset pairs. Next, PRECISE compares these independent PCs by computing the cosine similarity matrix and decomposing it by SVD [22]:

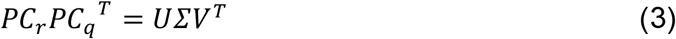

where *U* and *V* represent orthogonal (of size d) transformations on the reference and query PCs, respectively, and *Σ* is a diagonal matrix. *U* and *V* are then used to align the PCs, yielding the so-called Principal Vectors (PVs), such that:

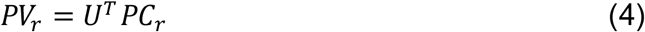

and

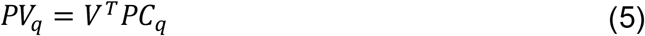

*PV_r_* and *PV_q_* are the principal vectors of the reference and the query datasets, respectively, retaining the same information as the principal components. However, these PVs have now a one-to-one correspondence as their cosine similarity matrix is diagonal (the matrix *Σ*). PVs are pairs of vectors 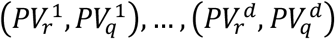 sorted in decreasing order based of similarity. To remove noisy components, we choose a limited number of PVs, *d*’, for further analysis, where the cosine similarity is higher than a certain threshold (0.3). The reference PVs, *PV_r_*, are then used to project and align both the scRNA-seq (reference) and the spatial transcriptomics (query) datasets:

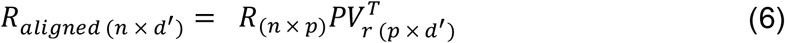

and

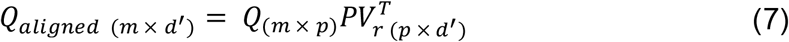

After aligning the datasets, SpaGE predicts the expression of the spatially unmeasured genes, *I = g -p*, from the scRNA-seq dataset. For each spatial cell *i ∈m*, we define the *k*-nearest-neighbors (*k* = 50) from the *n* dissociated scRNA-seq cells, using the cosine distance. Next, we calculate an array of weights *w_ij_* between spatial cell *i*. and its nearest neighbors *j ∈ NN*(ι). Out of the 50 neighbors, we only keep neighbors with positive cosine similarity with cell *i*. (i.e. cosine distance < 1), such that:

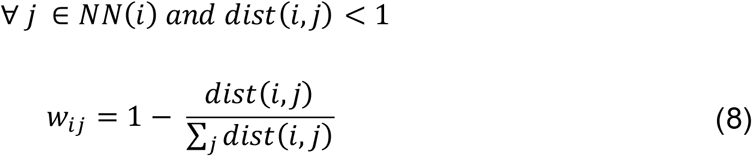

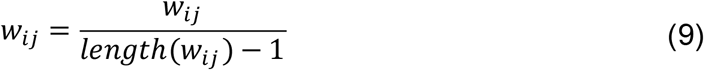

The predicted expression *Y_il_* of the set of spatially unmeasured genes *I* for cell *i*. is calculated as a weighted average of the nearest neighbors dissociated cells:

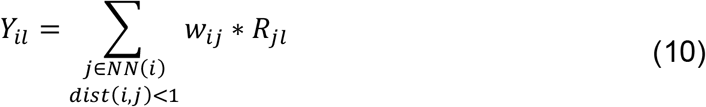

### Gene contribution to the integration

To evaluate the contribution of each gene in forming this common latent space *PV_r_*, we calculated the gene contribution *C_g_* of gene *g* as follows:

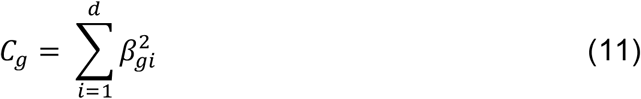

where *β_gi_* is the loading of gene *g* to the *i*-th principal vector in *PV_r_*, and *d* is the number of PVs in *PV_r_*. To obtain the top contributing genes, the *C_g_* values are sorted in descending order across all genes.

### Datasets

We used five dataset pairs (Table 1) composed of four scRNA-seq datasets (**AllenVISp** [19], **AllenSSp** [21], **Zeisel** [20], **Moffit** [4]) and three spatial transcriptomics datasets (**STARmap** [7], **osmFISH** [5], **MERFISH** [4]). The **AllenVISp** (GSE115746) and the **AllenSSp** datasets were downloaded from https://portal.brain-map.org/atlases-and-data/rnaseq. The **AllenVISp** is obtained from the ‘Cell Diversity in the Mouse Cortex – 2018’ release. The **AllenSSp** is obtained from the ‘Cell Diversity in the Mouse Cortex and Hippocampus’ release of October 2019. We downloaded the whole dataset and used the metadata to only select cells from the SSp region. The **Zeisel** dataset (GSE60361) was downloaded from http://linnarssonlab.org/cortex/, while the **Moffit** 10X dataset (GSE113576) was downloaded from GEO.

The **STARmap** dataset was downloaded from the STARmap resources website (https://www.starmapresources.com/data). We obtained the gene count matrix and the cell position information for the largest 1020-gene replicate. Cell locations and morphologies were identified using Python code provided by the original study (https://github.com/weallen/STARmap). The **osmFISH** dataset was downloaded as loom file from http://linnarssonlab.org/osmFISH/, we obtained the gene count matrix and the metadata using the loompy Python package. The **MERFISH** dataset was downloaded from Dryad repository (https://doi.org/10.5061/dryad.8t8s248), we used the first naïve female mouse (Animal_ID = 1).

### Data preprocessing

For all the scRNA-seq datasets, we filtered out genes expressed in less than 10 cells. No filtration was applied on the cells, except for the **AllenVISp** dataset for which we filtered low quality cells provided from the metadata (‘Low Quality’ and ‘No Class’ cells). For the **Zeisel** dataset, we only used the somatosensory cells excluding the hippocampus cells. Next, scRNA-seq datasets were normalized by dividing the counts within each cell by the total number of transcripts within that cell, scaling by 10^6^ and log(x+1) transformed. Further, we scaled the data by making each gene centered and scaled (zero mean and unit variance) using the SciPy Python package [23].

For spatial transcriptomics datasets all gene were used, except for the **MERFISH** dataset for which we removed the blanks genes and the *Fos* gene (non-numerical values). Additionally, we filtered out cells labeled as ‘Ambiguous’ from the **MERFISH** dataset. Similar to the Zeisel dataset, we only kept cells from cortical regions for the **osmFISH** dataset (‘Layer 2-3 lateral’, ‘Layer 2-3 medial’, ‘Layer 3-4’, ‘Layer 4’,’Layer 5’, ‘Layer 6’ and ‘Pia Layer 1’). No cells were filtered from the **STARmap** dataset. Further, each dataset was normalized by dividing the counts within each cell by the total number of transcripts within that cell, scaling by the median number of transcripts per cell, and log1p transformed. Similar to the scRNA-seq data, we scaled the spatial data using the SciPy Python package [23].

It is important to note that in all experiments, the scaled datasets are used as input for the alignment part, while the prediction is applied using the normalized version of the scRNA-seq dataset (Equation 10).

### Benchmarked methods

We compared the performance of SpaGE versus three state-of-the-art methods for data integration: Seurat, Liger, and gimVI. Seurat and Liger are available as R packages, while gimVI is available through the scVI Python package [24]. We were not able to include Harmony in the comparison, as the code to predict unmeasured gene expression is not available. During the benchmark, all methods were applied using their default settings, or the settings provided in the accompanying examples or vignettes. Data normalization and scaling were performed using the built-in functions in each package, *NormalizeData* and *ScaleData* functions in Seurat, *normalize* and *scaleNotCenter* functions in Liger, while gimVI implicitly preprocess the data while computing.

### Moran’s I statistic

The Moran’s I statistic [25] is a measure of spatial autocorrelation calculated for each spatial gene. The Moran’s I values can range from −1 to 1, where a value close to 1 indicates a clear spatial pattern, and a value close to 0 indicates random spatial expression, while a value close to −1 indicated a chess board like pattern. We calculated the Moran’s I using the following equation:

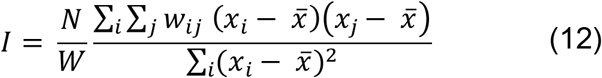

Where *x* is the gene expression array, 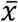 is the mean expression of gene *x, N* is the total number of spatial cells, *w_ij_* is a matrix containing spatial weights with zeros on the diagonal, and *W* is the sum of *w_ij_*. We calculated the spatial weights *w_ij_* using the XY coordinates of the spatial cells, for each cell we calculated the kNN using the spatial coordinates (k=4). We assigned *w_ij_* = 1 if *j* is in the nearest neighbors of *i*, otherwise *w_ij_* = 0.

### Down-sampling

For the 994 shared genes in the **STARmap_AllenVISp** dataset pair, we first selected the top 50 spatial genes with high Moran’s I statistic values to be used as test set. For the remaining 944 genes, we calculated the pairwise Pearson correlation between using the scRNA-seq dataset. If the absolute value of the correlation of two genes is larger than 0.7, we removed the gene with the lower variance. After removing highly correlated genes, we sorted the remaining genes according to their expression variance in the scRNA-seq dataset. We selected the top 10, 30, 50, 100 and 200 genes with high variance, these genes were used for alignment of the two datasets and prediction of the expression of the test genes. The prediction performance of these gene sets was compared with using all 944 genes.

## Availability of data and materials

The implementation code of SpaGE, as well as the benchmarking code, is available in the GitHub repository, at https://github.com/tabdelaal/SpaGE. The code is released under MIT license. All datasets used are publicly available data.

