## Supplementary Figures for "SpaGE: Spatial Gene Enhancement using scRNA-seq"

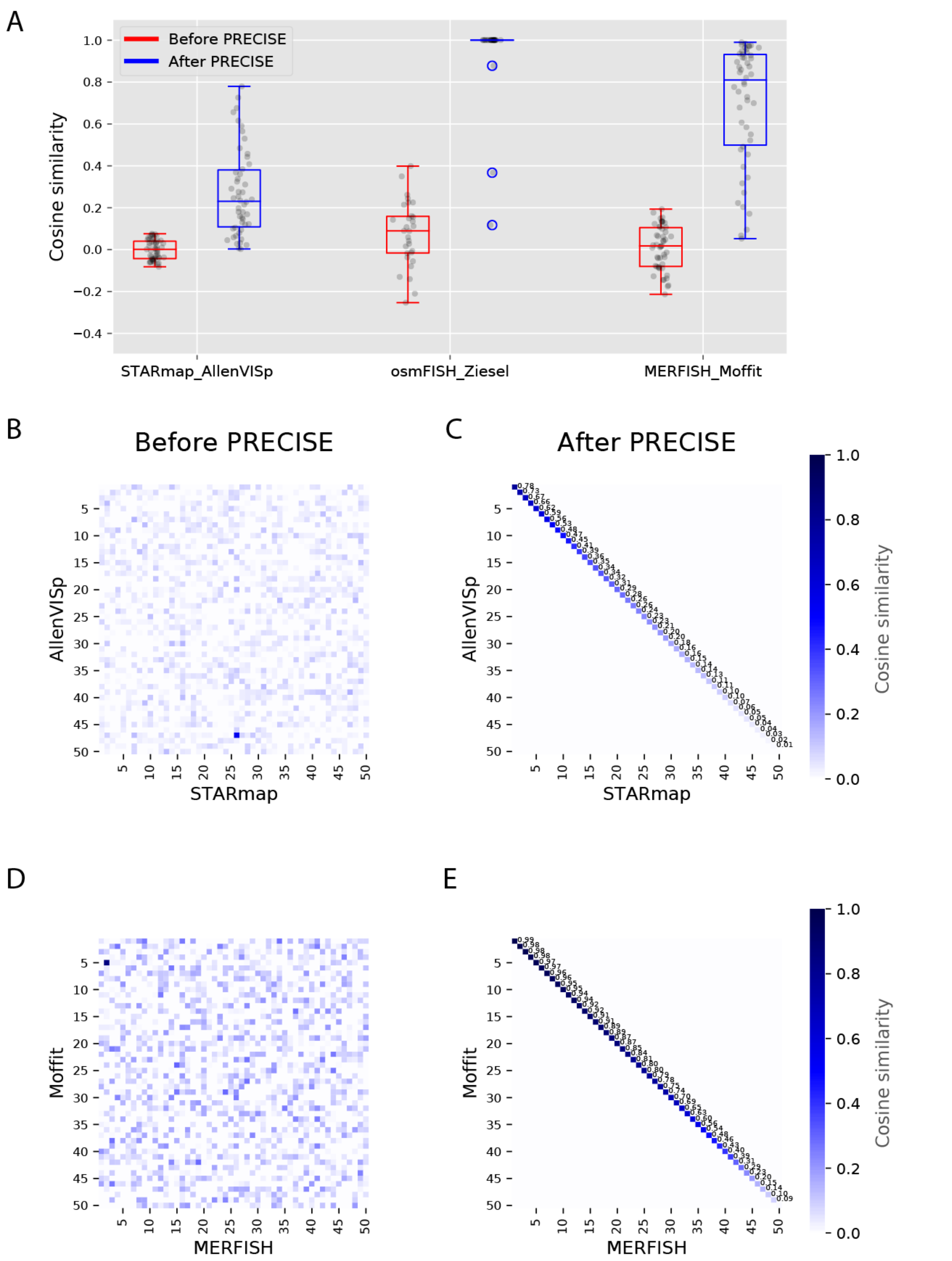


**Figure S1 Pairwise cosine similarity matrices before and after PRECISE. (A)** Boxplots showing the diagonal (one-to-one) Cosine similarity between the independent *PCs* (before PRECISE) of both datasets in each dataset pair, and between the *PVs* (after PRECISE). **(B,D)** Pairwise cosine similarity matrices between the *PCs* (before PRECISE) of the **(B) STARmap_AllenVISp** and the **(D) MERFISH_Moffit** dataset pairs, showing no one-to-one correspondence. **(C,E)** Pairwise cosine similarity matrices between the *PVs* (after PRECISE) of the **(C) STARmap_AllenVISp** and the **(E) MERFISH_Moffit** dataset pairs, showing a clear one-to-one diagonal similarity.

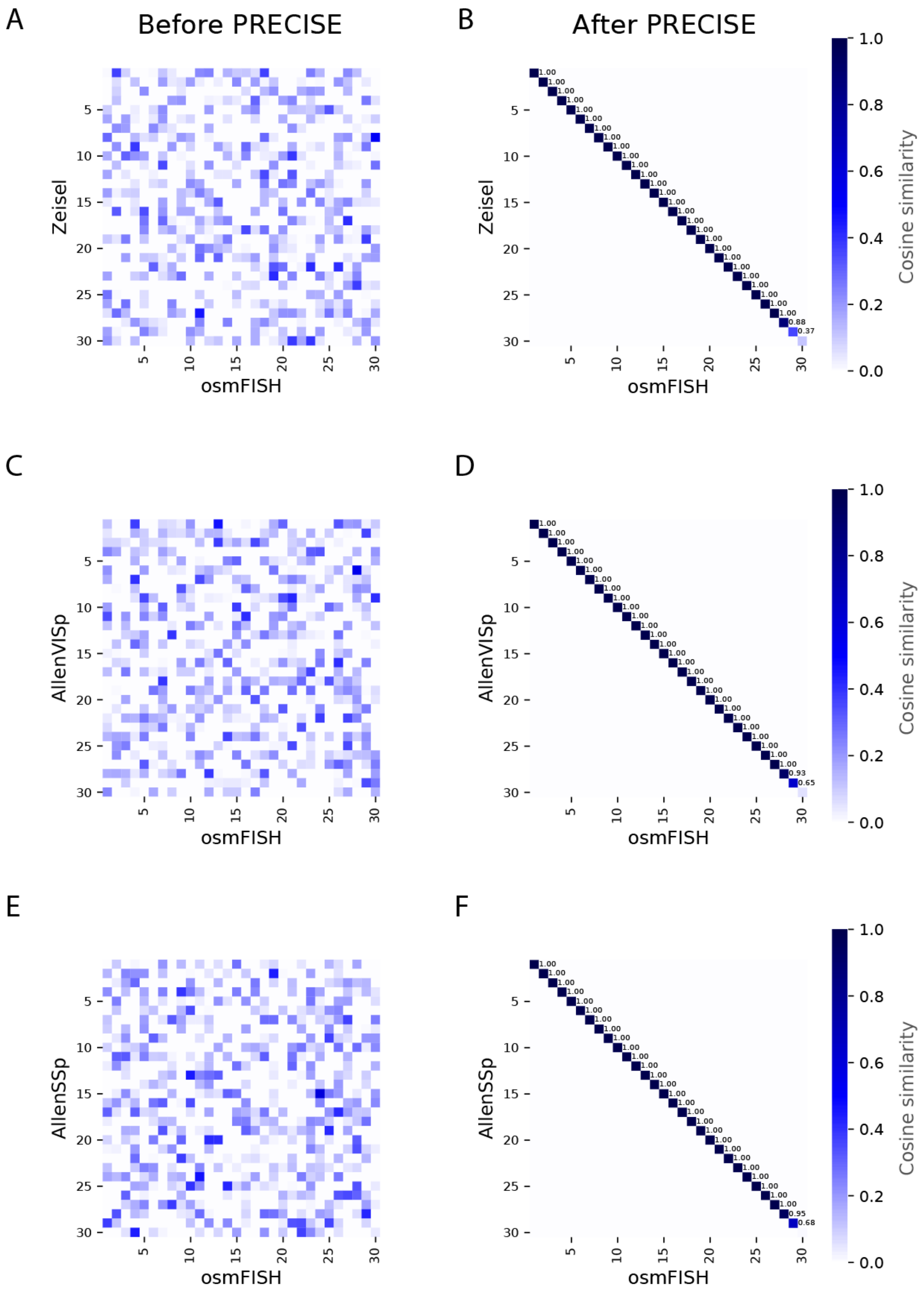


**Figure S2 Pairwise cosine similarity matrices before and after PRECISE. (A,C,E)** Pairwise cosine similarity matrices between the *PCs* (before PRECISE) of the **(A) osmFISH_Zeisel**, the **(C) osmFISH_AllenVISp** and the **(E) osmFISH_AllenSSp** dataset pairs, showing no one-to-one correspondence. **(B,D,F)** Pairwise cosine similarity matrices between the *PVs* (after PRECISE) of the **(B) osmFISH_Zeisel**, the **(D) osmFISH_AllenVISp** and the **(F) osmFISH_AllenSSp** dataset pairs, showing a clear one-to-one diagonal similarity.

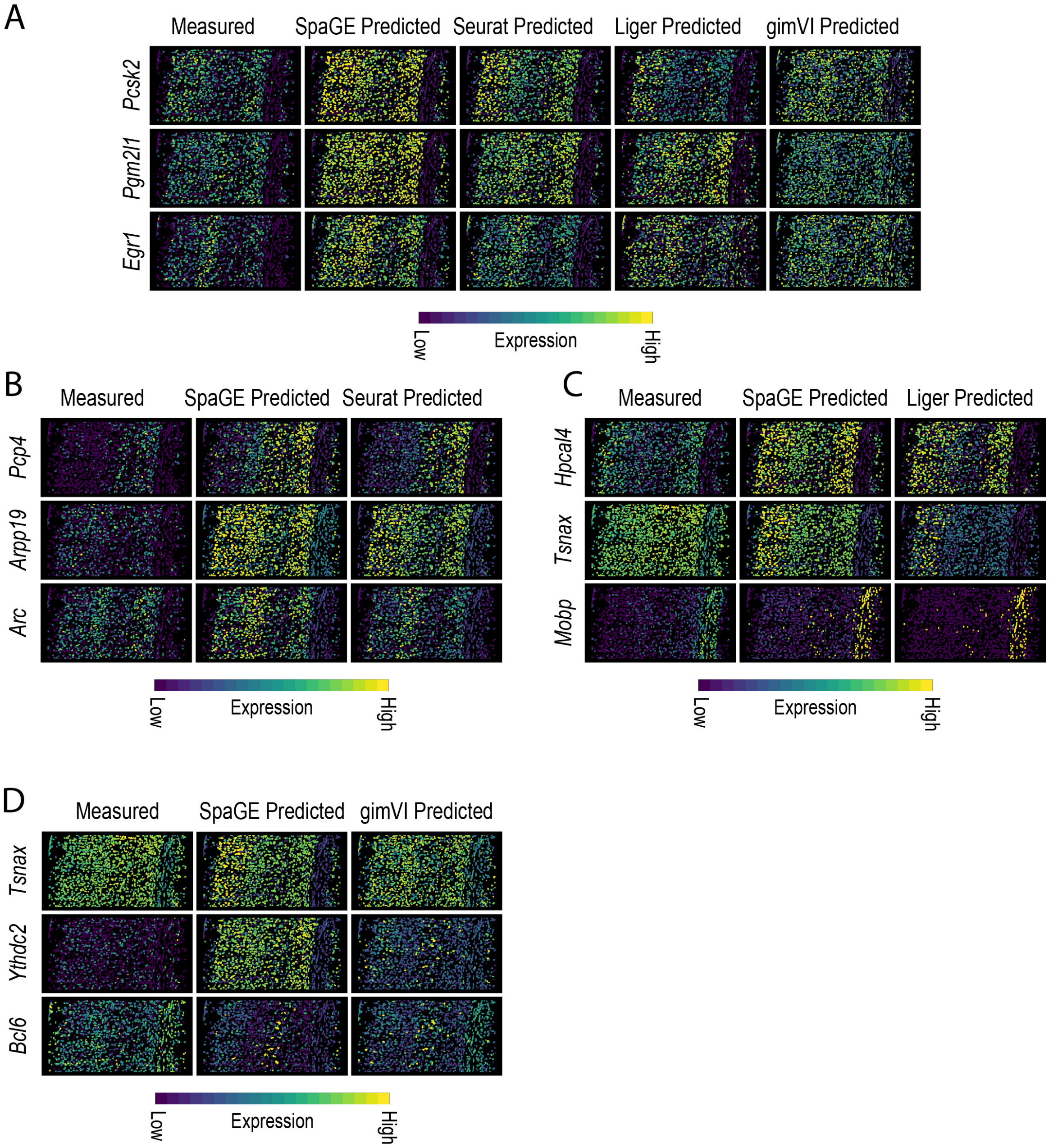


**Figure S3 Top predicted genes of each method using the STARmap_AllenVISp dataset pair. (A)** Comparison of the top 3 genes predicted by SpaGE. Each row corresponds to a single gene, first column from the left shows the measured spatial gene expression in the **STARmap** dataset, while other columns show the corresponding predicted expression pattern by SpaGE, Seurat, Liger and gimVI. **(B-D)** Comparison of the top 3 genes predicted by **(B)** Seurat, **(C)** Liger and **(D)** gimVI, excluding the top 10 predicted genes by SpaGE. Each row corresponds to a single gene, first column from the left shows the measured spatial gene expression in the **STARmap** dataset, the middle column shows the corresponding predicted expression pattern by SpaGE, while the right column shows predicted expression pattern by **(B)** Seurat, **(C)** Liger and **(D)** gimVI. All predictions were obtained using the leave-one-gene-out cross validation experiment.

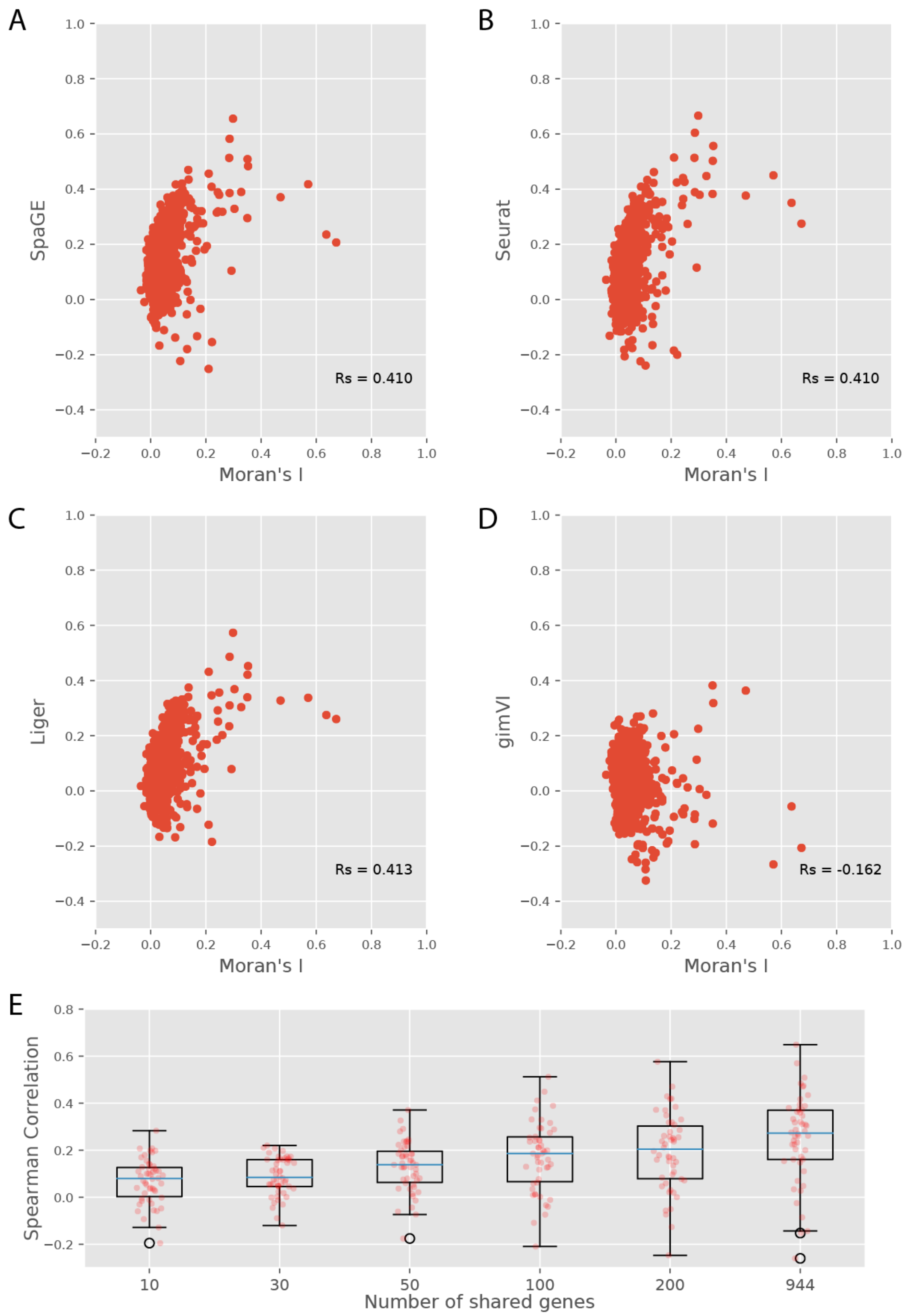


**Figure S4 (A-D)** Scatter plots showing the relation between the Moran's I statistic and the prediction correlation of each gene, using the **STARmap_AllenVISp** dataset pair. Moran's I (x-axis) are calculated using the **STARmap** dataset and prediction correlation values (y-axis) were obtained by **(A)** SpaGE, **(B)** Seurat, **(C)** Liger and **(D)** gimVI. The Rs values correspond to the Spearman Rank correlation between the Moran's I statistic and the prediction performance of each method. **(E)** Boxplots showing the prediction performance of a test set of 50 genes, in terms of Spearman Rank correlations, using downsampled sets of 10, 30, 50, 100 and 200 shared genes compared to using all 944 genes in the **STARmap_AllenVISp** dataset pair.

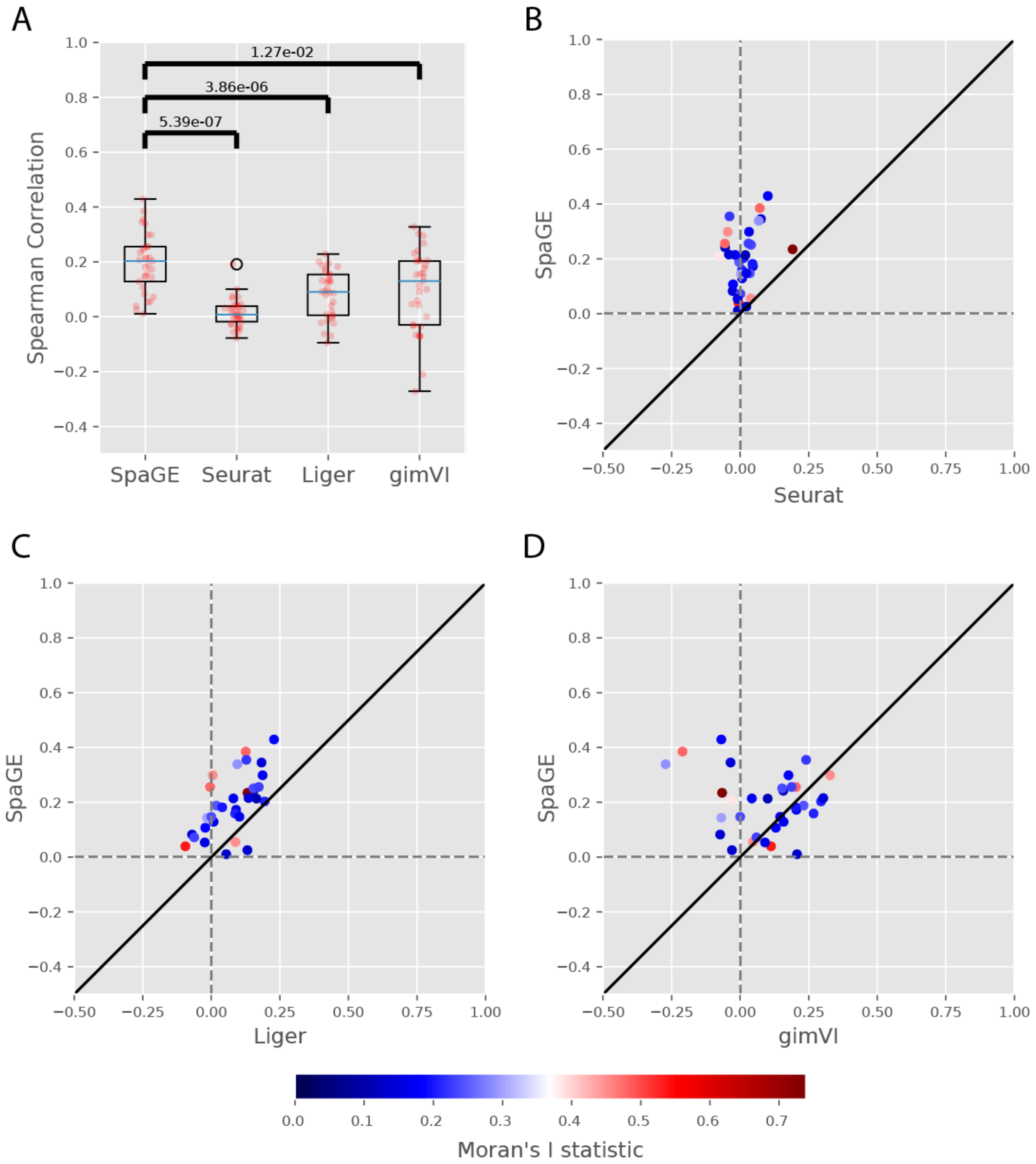


**Figure S5 Prediction performance comparison for the osmFISH_Zeisel dataset pair. (A)** Boxplots showing the Spearman correlations for the leave-one-gene-out cross validation experiment for each method. The blue lines show the median correlation across all genes with a better performance for SpaGE. The red dots show the correlation values for individual genes. The p-values show the significant difference between all correlation values of SpaGE and each other method, using a paired Wilcoxon rank-sum test. **(B-D)** Detailed performance comparison between SpaGE and **(B**) Seurat, **(C)** Liger, **(D)** gimVI. These scatter plots show the correlation value of each gene across two methods. The solid black line is the y=x line, the dashed lines show the zero correlation. Points are colored according to the Moran's I statistic of each gene. All scatter plots show that the majority of the genes are skewed above the y=x line, showing an overall better performance of SpaGE over other methods.

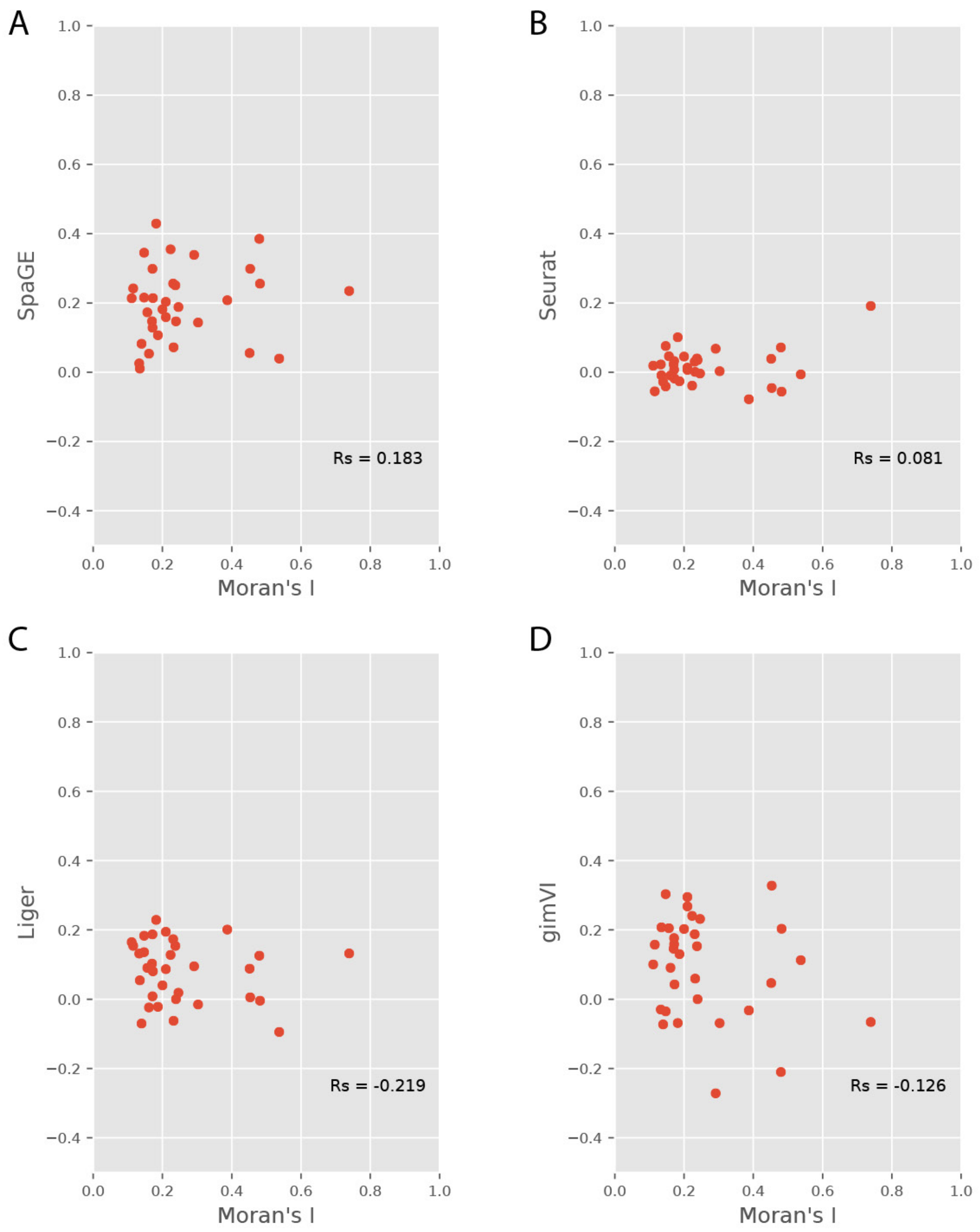


**Figure S6 (A-D)** Scatter plots showing the relation between the Moran's I statistic and the prediction correlation of each gene, using the **osmFISH_Zeisel** dataset pair. Moran's I (x-axis) are calculated using the **osmFISH** dataset and prediction correlation values (y-axis) were obtained by **(A)** SpaGE, **(B)** Seurat, **(C)** Liger and **(D)** gimVI. The Rs values correspond to the Spearman Rank correlation between the Moran's I statistic and the prediction performance of each method.

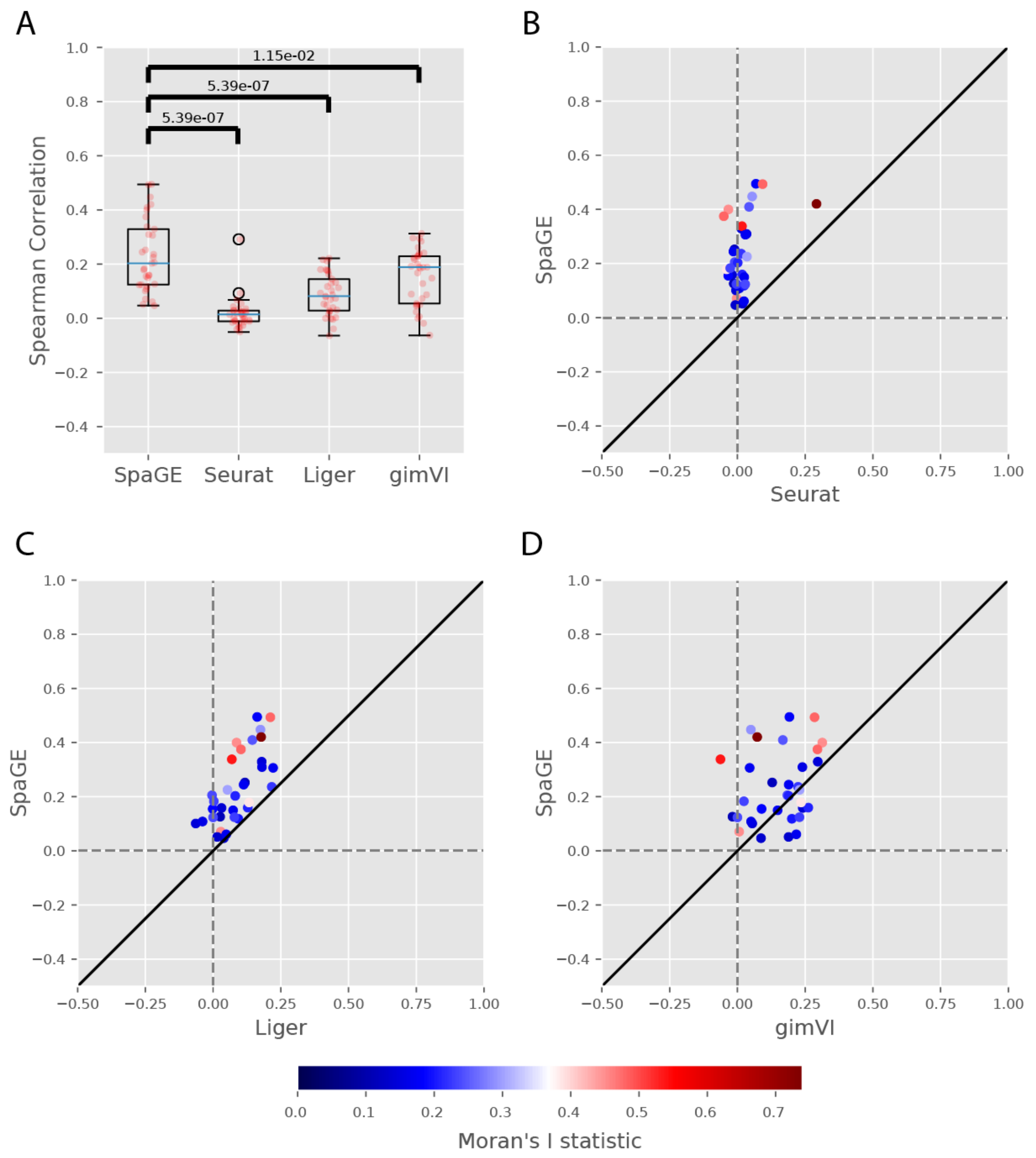


**Figure S7 Prediction performance comparison for the osmFISH_AllenVISp dataset pair. (A)** Boxplots showing the Spearman correlations for the leave-one-gene-out cross validation experiment for each method. The blue lines show the median correlation across all genes with a better performance for SpaGE. The red dots show the correlation values for individual genes. The p-values show the significant difference between all correlation values of SpaGE and each other method, using a paired Wilcoxon rank-sum test. **(B-D)** Detailed performance comparison between SpaGE and **(B)** Seurat, **(C)** Liger, **(D)** gimVI. These scatter plots show the correlation value of each gene across two methods. The solid black line is the y=x line, the dashed lines show the zero correlation. Points are colored according to the Moran's I statistic of each gene. All scatter plots show that the majority of the genes are skewed above the y=x line, showing an overall better performance of SpaGE over other methods.

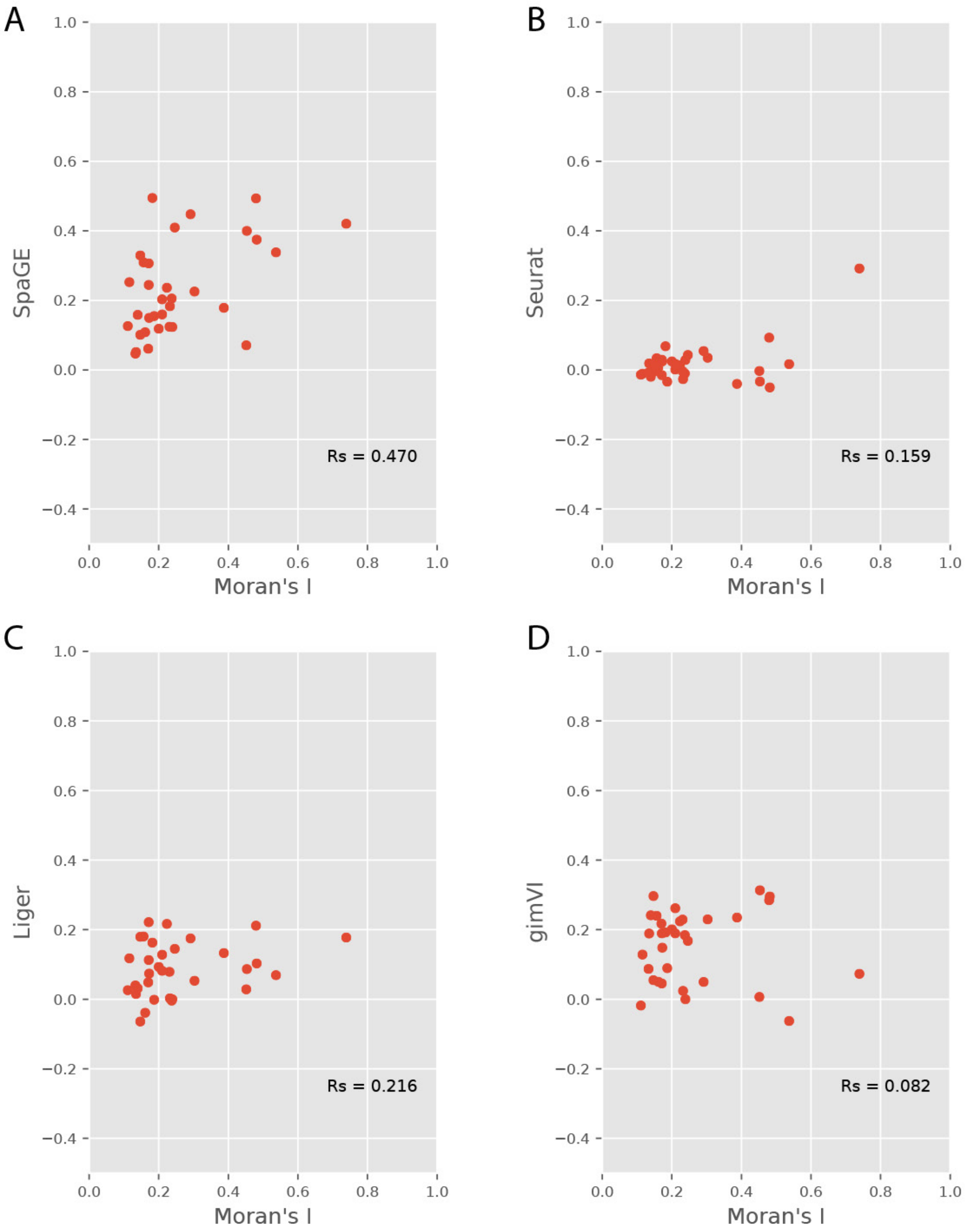


**Figure S8 (A-D)** Scatter plots showing the relation between the Moran's I statistic and the prediction correlation of each gene, using the **osmFISH_AllenVISp** dataset pair. Moran's I (x-axis) are calculated using the **osmFISH** dataset and prediction correlation values (y-axis) were obtained by **(A)** SpaGE, **(B)** Seurat, **(C)** Liger and **(D)** gimVI. The Rs values correspond to the Spearman Rank correlation between the Moran's I statistic and the prediction performance of each method.

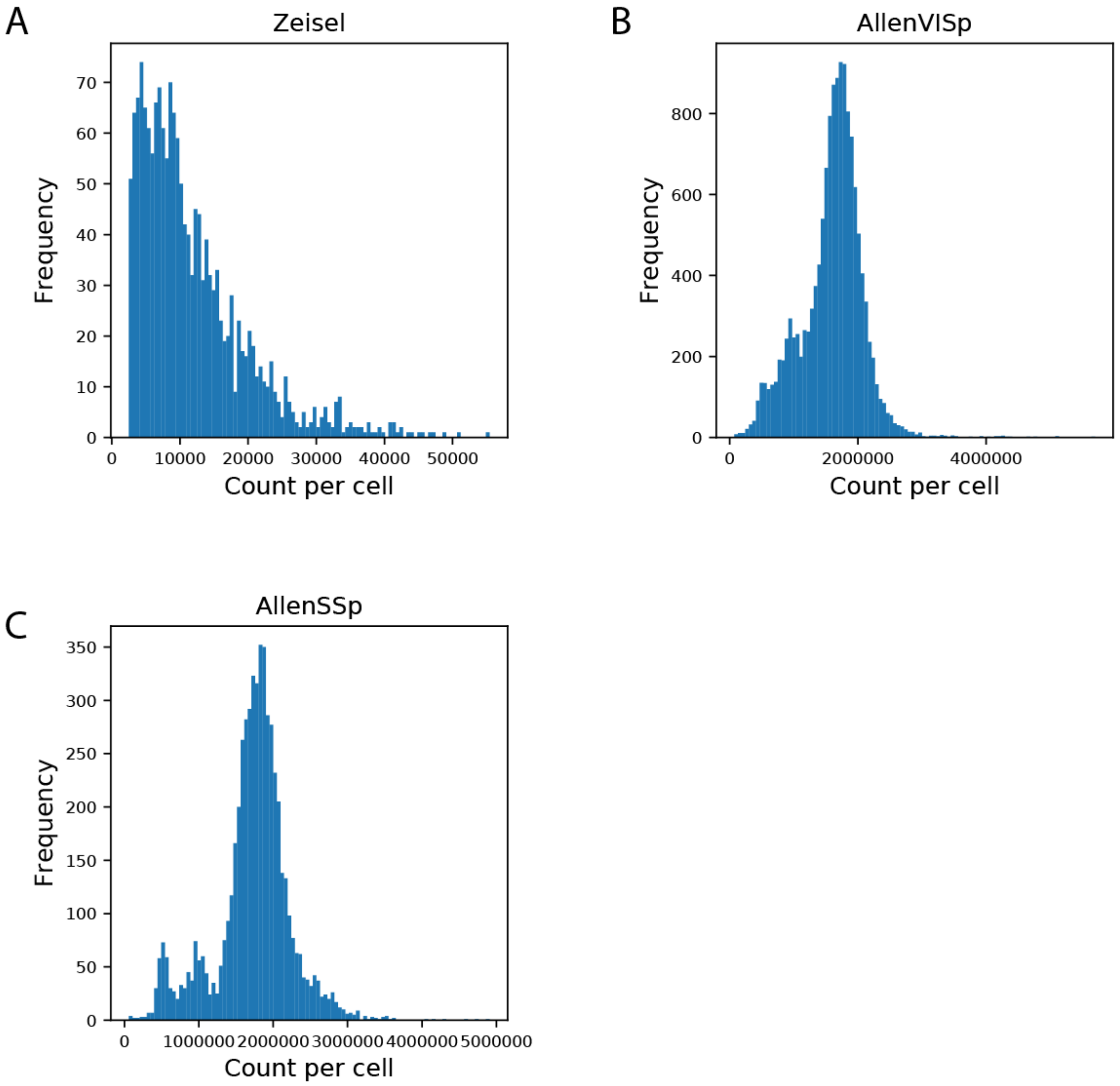


**Figure S9 Sequencing depth across difference scRNA-seq reference datasets.** Histograms showing the distribution of the RNA count per cell of the **(A) Zeisel**, **(B) AllenVISp**, and **(C) AllenSSp** datasets, respectively. The **AllenVISp** and **AllenSSp** datasets have comparable sequencing depths, while the sequencing depth of the **Zeisel** dataset is much lower.

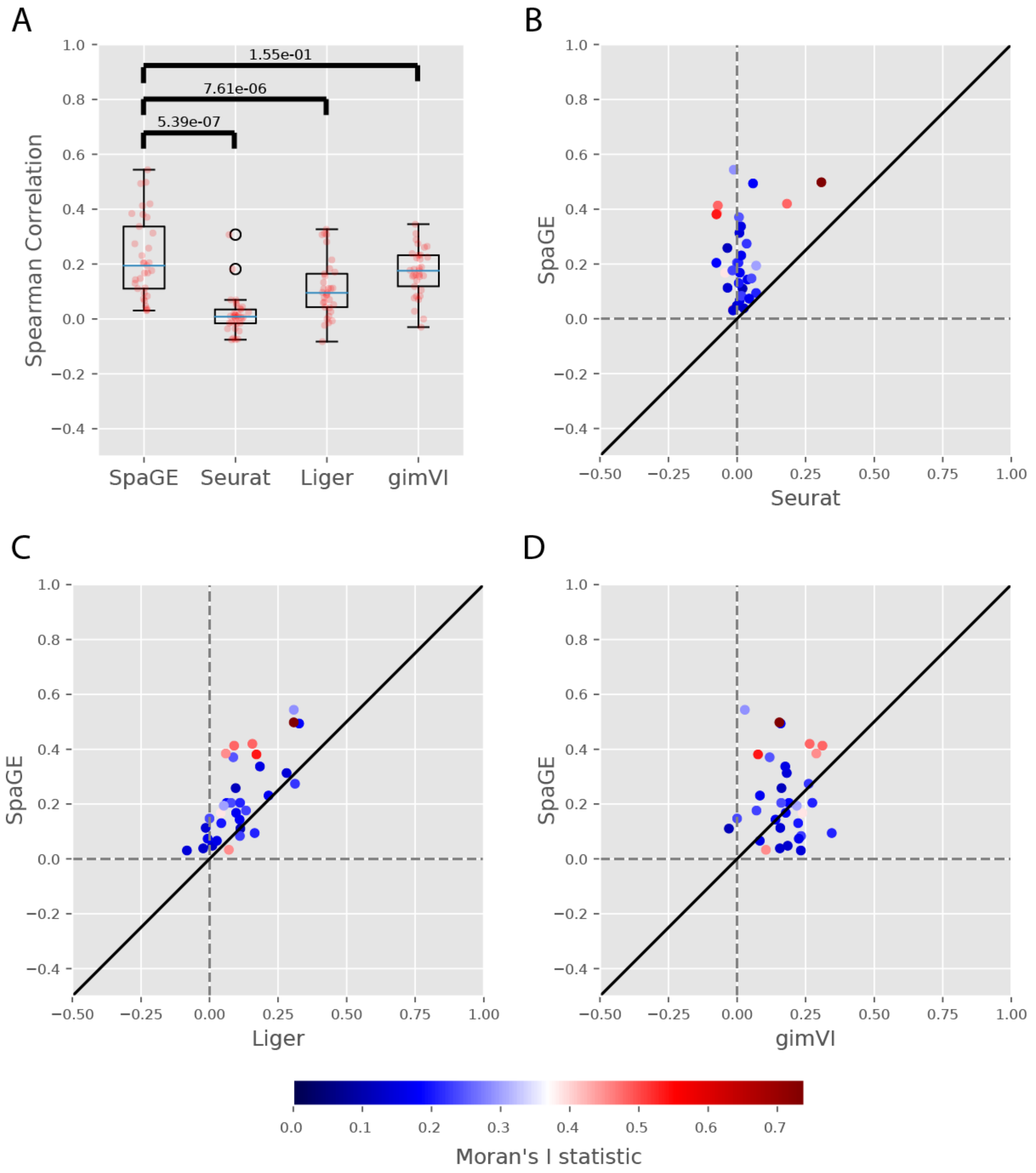


**Figure S10 Prediction performance comparison for the osmFISH_AllenSSp dataset pair. (A)** Boxplots showing the Spearman correlations for the leave-one-gene-out cross validation experiment for each method. The blue lines show the median correlation across all genes with a better performance for SpaGE. The red dots show the correlation values for individual genes. The p-values show the significant difference between all correlation values of SpaGE and each other method, using a paired Wilcoxon rank-sum test. **(B-D)** Detailed performance comparison between SpaGE and **(B)** Seurat, **(C)** Liger, **(D)** gimVI. These scatter plots show the correlation value of each gene across two methods. The solid black line is the y=x line, the dashed lines show the zero correlation. Points are colored according to the Moran's I statistic of each gene. All scatter plots show that the majority of the genes are skewed above the y=x line, showing an overall better performance of SpaGE over other methods.

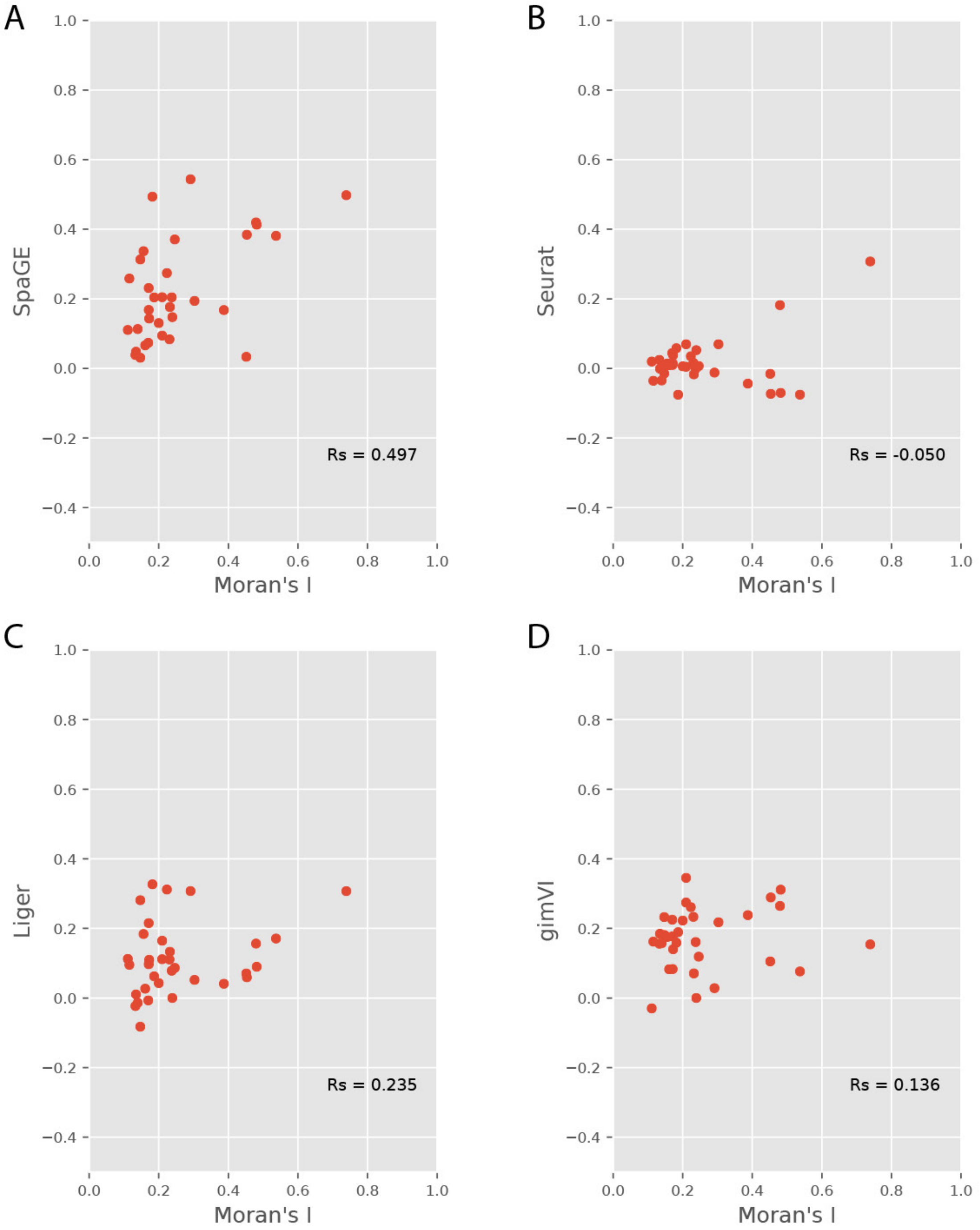


**Figure S11 (A-D)** Scatter plots showing the relation between the Moran's I statistic and the prediction correlation of each gene, using the **osmFISH_AllenSSp** dataset pair. Moran's I (x-axis) are calculated using the **osmFISH** dataset and prediction correlation values (y-axis) were obtained by **(A)** SpaGE, **(B)** Seurat, **(C)** Liger and **(D)** gimVI. The Rs values correspond to the Spearman Rank correlation between the Moran's I statistic and the prediction performance of each method.
